# A telomere to telomere phased genome assembly and annotation for the Australian central bearded dragon *Pogona vitticeps*

**DOI:** 10.1101/2025.05.01.651798

**Authors:** Hardip R. Patel, Kirat Alreja, Andre L.M. Reis, J King Chang, Zahra A. Chew, Hyungtaek Jung, Jillian M. Hammond, Ira W. Deveson, Aurora Ruiz-Herrera, Laia Marin-Gual, Clare E. Holleley, Xiuwen Zhang, Nicholas C. Lister, Sarah Whiteley, Lei Xiong, Duminda S.B. Dissanayake, Paul D. Waters, Arthur Georges

## Abstract

**Background:** The central bearded dragon (*Pogona vitticeps)* is widely distributed in central eastern Australia and adapts readily to captivity. Among other attributes, it is distinctive because it undergoes sex reversal from ZZ genotypic males to phenotypic females at high incubation temperatures. Here, we report an annotated telomere to telomere phased assembly of the genome of a female ZW central bearded dragon.

**Results:** Genome assembly length is 1.75 Gbp with a scaffold N50 of 266.2 Mbp, N90 of 28.1 Mbp, 26 gaps and 42.2% GC content. Most (99.6%) of the reference assembly is scaffolded into 6 macrochromosomes and 10 microchromosomes, including the Z and W microchromosomes, corresponding to the karyotype. The genome assembly exceeds standard recommended by the Earth Biogenome Project (6CQ40): 0.003% collapsed sequence, 0.03% false expansions, 99.8% k-mer completeness, 97.9% complete single copy BUSCO genes and an average of 93.5% of transcriptome data mappable back to the genome assembly. The mitochondrial genome (16,731 bp) and the model rDNA repeat unit (length 9.5 Kbp) were assembled. Male vertebrate sex genes *Amh* and *Amhr2* were discovered as copies in the small non-recombining region of the Z chromosome, absent from the W chromosome.

This, coupled with the prior discovery of differential Z and W transcriptional isoform composition arising from pseudoautosomal sex gene *Nr5a1*, suggests that complex interactions between these genes, their autosomal copies and their resultant transcription factors and intermediaries, determines sex in the bearded dragon.

**Conclusion:** This high-quality assembly will serve as a resource to enable and accelerate research into the unusual reproductive attributes of this species and for comparative studies across the Agamidae and reptiles more generally.

**Species Taxonomy:** Eukaryota; Animalia; Chordata; Reptilia; Squamata; Iguania; Agamidae; Amphibolurinae; *Pogona*; *Pogona vitticeps* (Ahl, 1926) (NCBI:txid103695).

**Graphical Abstract:** 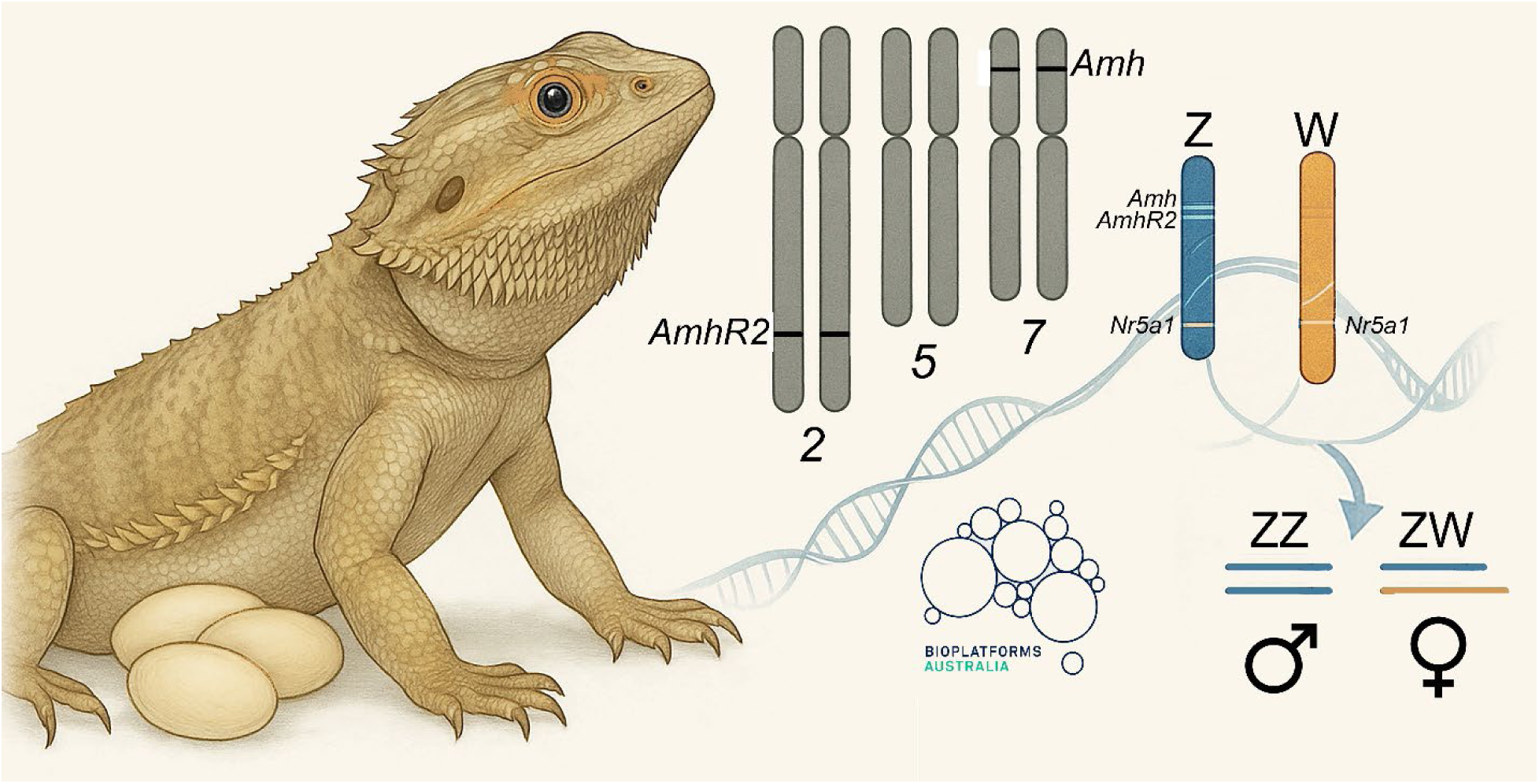

## Introduction

The family Agamidae, commonly known as dragon lizards, is a diverse group of lizards found in Africa, Asia, Australia, the Western Pacific, and warmer regions of Southern Europe. The Agamidae family is well represented in Australia, in part because of their successful radiation in response to the progressive aridification of the Australian continent during the Pleistocene. New species are continually being described, but on recent count they comprise 81 species in 15 genera (Cogger 2018) that occupy a very wide array of habitats ranging from the inland deserts to the mesic habitats of the coast and the Australian Alps below the tree-line. The family includes some iconic species such as the thorny devil *Moloch horridus* and the frillneck lizard *Chlamydosaurus kingii*. Less spectacular perhaps is the central bearded dragon *Pogona vitticeps* (Ahl, 1926), a widely distributed species of Amphibolurine dragon common in central eastern Australia (Figure 1). The bearded dragon feeds on insects and other invertebrates, but a substantial component of the diet of adults is vegetable matter. It lives in the dry sclerophyll forests and woodlands in the southeast of its range, mallee and arid acacia scrublands further north and west, and the sandy deserts of the interior. Semi-arboreal, the species often perches on fallen timber and tree branches only to retreat to ground cover when disturbed.

**Figure 1.**
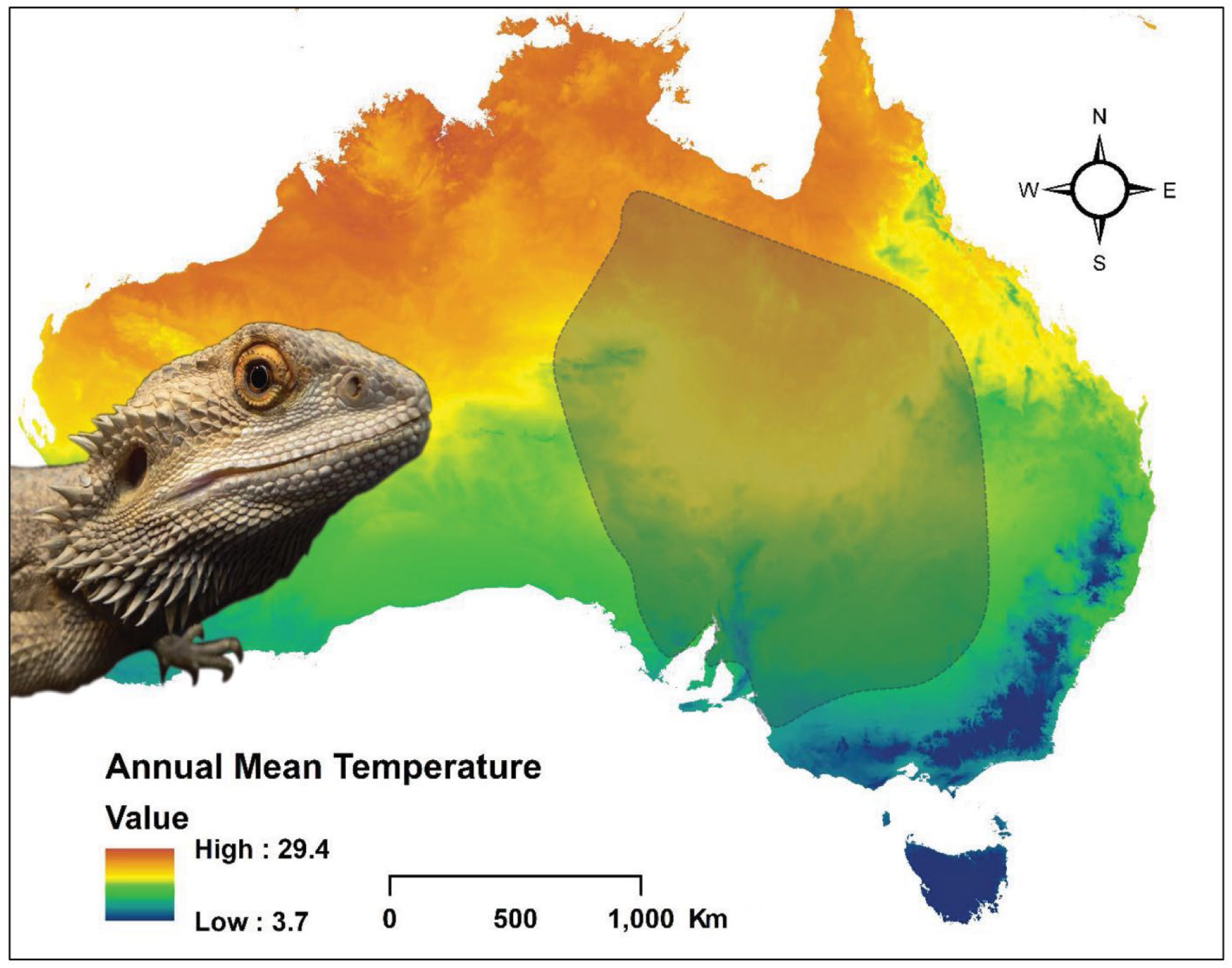
The central bearded dragon *Pogona vitticeps* and the distribution of the species based on records from Australian museums (via Atlas of Living Australia https://www.ala.org.au/).

Central bearded dragons adapt readily to captivity, lay large clutches of eggs several times per season, and are commonly kept as a pet in Europe, Asia, and North America. These attributes also increase its value as a popular reptile research model in a range of disciplines (Ollonen et al., 2018; Bonnan et al., 2024; Chandrasekara et al., 2024; Fenk et al., 2024; Nagashima et al., 2024; Razmadze et al., 2024). Central bearded dragons are a particularly compelling model species for sex determination because they display temperature-induced sex reversal in the laboratory and in the wild (Quinn et al., 2007; Holleley et al., 2015; Castelli et al., 2021). The sex chromosomes of central bearded dragons are poorly differentiated morphologically. They exhibit female heterogamety (ZZ/ZW sex chromosome system, Ezaz et al., 2005) with 6 macrochromosome pairs and 10 microchromosome pairs (Witten, 1983) that includes the sex microchromosome pair (Ezaz et al., 2005). BAC sequences have been physically mapped uniquely to each of the chromosomes (Young et al., 2013; Deakin et al., 2016).

Sex determination in this species is particularly subtle until now with no substantial difference between the Z and W chromosome gene content or single-copy sequence (Zhang et al., 2022). The developmental program initiated by chromosomal sex determination can be reversed by high incubation temperature, allowing for investigations of environmental influences on fundamental developmental processes. Research in these areas of interest will be greatly facilitated by applying modern sequencing technologies to generate a high-quality draft genome assembly for the central bearded dragon. The ability to generate telomere to telomere (T2T) assemblies of the sex chromosomes and identify the non-recombining regions within which lies any master sex determining gene will greatly narrow the field of candidate sex determining genes in species with chromosomal sex determination. Furthermore, the disaggregation of the Z and W sex chromosome haplotypes (phasing) will allow comparisons of the Z and W sequences to gauge putative loss or difference in function of key sex gene candidates.

In this paper, we present a draft annotated telomere to telomere phased assembly of the genome of the Australian central bearded dragon as a resource to enable and accelerate research into the unusual reproductive attributes of this species and for comparative studies across the Agamidae and reptiles more generally. This is a vastly improved assembly in comparison with an earlier assembly based on Illumina short-read technology published in 2015 (Georges et al., 2015).

## Materials and Methods

### Sample collection

DNA samples were obtained from a blood sample taken from a single female *Pogona vitticeps* (RadMum, UCID Pit_001003342236) collected on 15-Mar-2011 on a road verge 62 km west of Eulo on Adventure Way, Queensland (GPS −28.099000 144.433000). It was verified as a ZW female using sex-linked polymerase chain reaction (PCR) markers (Holleley et al., 2015).

An additional 3 adult individuals were sampled to provide tissues (brain, heart, kidney, liver, lung, skeletal muscle, testes, ovary), complemented by embryonic brain and gonad, for transcriptomics (Table S2).

### Extraction and Sequencing

We generated sequencing data using three platforms – PacBio HiFi, ONT ultralong reads and HiC generated using the Arima Genomics protocols (Figure 2). Illumina short read DNA data were previously generated (Georges et al., 2015). Transcriptome data were generated using the Illumina platform. All sequence data generated in this study are available from NCBI SRA under BioProject ID PRJNA1252275.

**Figure 2.**
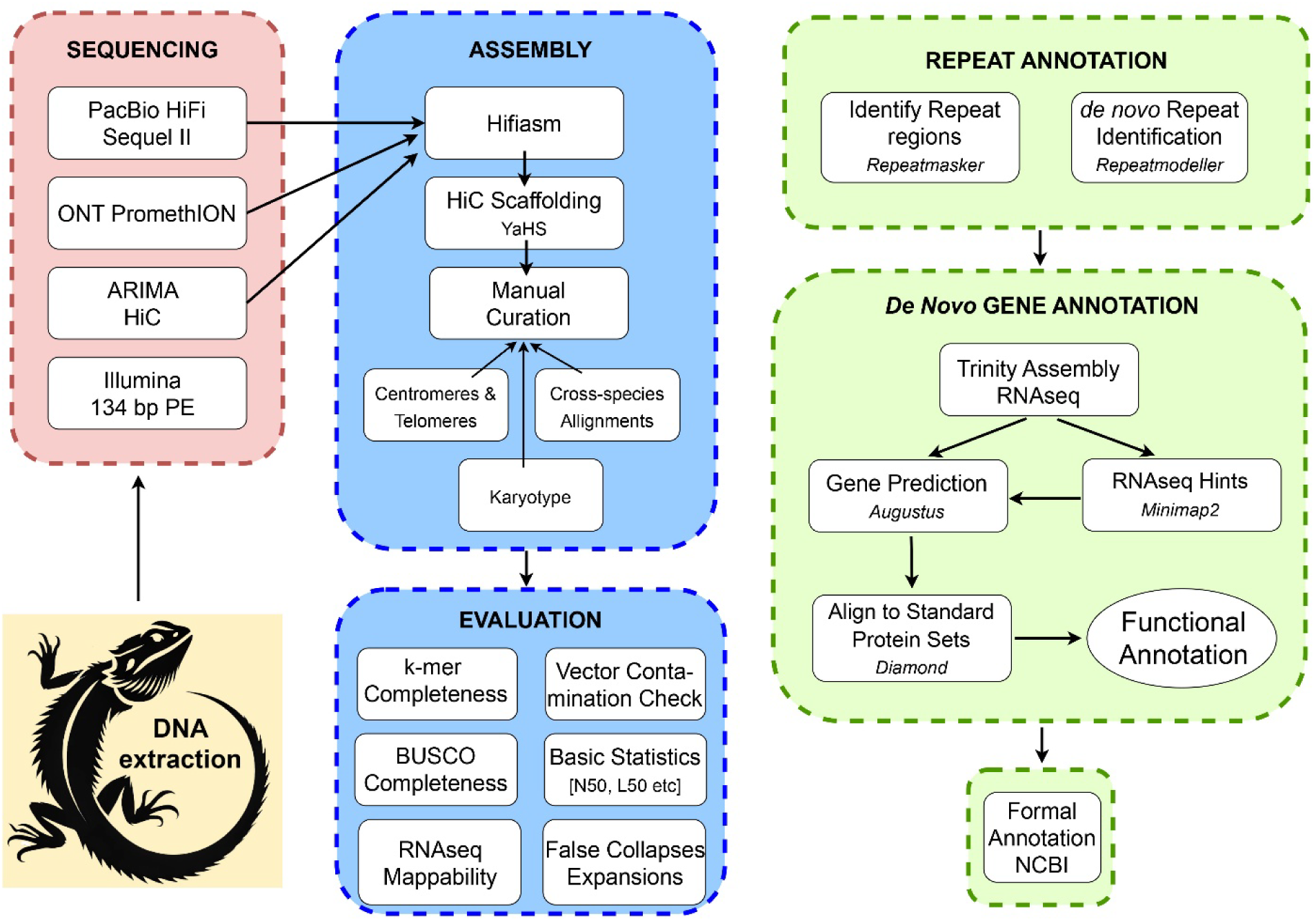
Schematic overview of workflow for sequencing, assembly and annotation of the genome of the central bearded dragon *Pogona vitticeps.* Target: Earth Biogenomes Project standard 6CQ40 (Lawniczak et al., 2022). Illumina 134 bp PE reads (Table S6) were not used directly in the assembly, but for quality assessment of the genome. Quality control workflow not shown. Repeat annotation was undertaken with Repeatmasker (4.1.2-p1, Smit et al., 2013-2015). Refer to Table S1 for software used in this project.

#### PacBio HiFi

Genomic DNA was extracted from blood of the focal ZW individual by PacBio Asia (Singapore) and sequenced using two flow cells on a PacBio Sequel II (Table S3). HiFi data were processed using *cutadapt* (v3.7, parameters: --anywhere --error-rate 0.1 --overlap 25 --match-read-wildcards --revcomp --discard-trimmed) to remove reads containing PacBio primers and adaptor sequences. This step removes putative chimeric sequences.

#### ONT PromethION

Genomic DNA was extracted from blood of the focal ZW individual (Table S4) using the salting out procedure (Miller et al., 1988) and spooled to enrich for high molecular weight DNA. DNA was shipped to the Garvan Institute of Medical Research in Sydney. Library preparation was performed with 3 µg of DNA as input, using the SQK-LSK109 kit (Oxford Nanopore Technologies, UK) and sequenced across 4 x promethION (FLO-PRO002) flow cells, with washes (EXP-WSH004) performed when sequencing dropped.

A second extraction was performed on 10 µl of blood using the Circulomics UHMW extraction kit, following the “Nucleated blood” protocol, obtaining approximately 60 µg of ultra-high molecular weight DNA. Library preparation was then performed using a pre-release version of the SQK-ULK001 kit from Oxford Nanopore Technologies, which uses the RAP adapter. The library was then loaded onto one promethION (FLO-PRO002) flow cell with washes (EXP-WSH004) performed at 24 and 48 hours to increase output.

ONT basecalling was performed using the *buttery-eel* (v0.4.2+dorado7.2.13, parameters: --config dna_r9.4.1_450bps_hac_prom.cfg --detect_mid_strand_adapter --trim_adapters --detect_adapter --do_read_splitting --qscore 7). Parameters were chosen to remove reads with average quality value score <7, remove adapters at 5’ or 3’ ends of sequence, and to split reads if adapters were in the middle of the read.

#### HiC

A blood sample from the focal ZW individual (Table S5) was used for HiC. Blood sample was processed by the Biomolecular Resource Facility (BRF) at the Australian National University using the Arima HiC 2.0 kit for library preparation and sequencing with two flow cells on an Illumina NovaSeq 6000 NovaSeq 6000, S1 300 cycles kit 2×150 bp.

#### RNA

Total RNA was extracted from adult brain, liver, heart and ovary/testis (Table S2) by the Garvan Institute of Medical Research (Sydney). Tissue extracts were homogenized using T10 Basic ULTRA-TURRAX® Homogenizer (IKA, Staufen im Breisgau, Germany) and extracted using TRIzol reagent following the manufacturer’s instructions, purifying with an isopropanol precipitation. Seventy-five bp single-end reads were generated for recent samples on the Illumina NextSeq 500 platform at the Ramaciotti Centre for Genomics (UNSW, Sydney, Australia). Some earlier samples generated 100 bp PE reads.

RNAseq from three embryonic gonads were sourced from Whiteley *et al*. (2022) and RNAseq from three embyronic brains were sourced from Whiteley *et al.,* (2021) and Wagner *et al.,* (2023) (Table S2).

### Assembly

All data analyses were performed on the high-performance computing facility, Gadi, hosted by Australia’s National Computational Infrastructure (NCI, https://nci.org.au). Scripts are available at https://github.com/kango2/ausarg.

#### Primary genome assembly

PacBio HiFi, ONT and HiC sequence data were used to generate interim haplotype assemblies and an interim pseudohaplotype (=consensus haplotype) assembly using *hifiasm* (v0.19.8, Cheng et al. 2021, 2022, default parameters). HiC data were aligned to the interim pseudohaplotype and haplotype assembly using the *Arima Genomics alignment pipeline* (v03, https://github.com/ArimaGenomics/mapping_pipeline, last accessed 16-Apr-2025) following the user guide for scaffolding and assessing the accuracy of assembly. HiC read alignments were processed using *YaHS* (v1.1, Zhou et al. 2022, parameters: -r 10000, 20000, 50000, 100000, 200000, 500000, 1000000, 1500000 --no-contig-ec -e GATC,GANTC,CTNAG,TTAA) to generate scaffolds. Range resolution parameter (-r) in *YaHS* was restricted to 1500000 to ensure separation of microchromosomes into individual scaffolds. Contig correction was disabled to maintain the original contig structure produced by *hifiasm*.

HiC contact maps were processed and visualised using *Juicer (*v1.5, Durand et al., 2016). Read depth, GC content, and telomere locations for *YaHS* scaffolds >1 Mbp length were visually inspected. One scaffold in the pseudohaplotype assembly contained internal telomeric repeat and contact pattern of a mis-join, owing to incorrect contig assembly by *hifiasm*. Similar error was observed for one scaffold in haplotype 2 as a result of scaffolding error (Figure S2). YaHS was rerun without --no-contig-ec parameter, which is the default behaviour that fixed these errors.

#### Reference genome assembly

The karyotype was obtained from Witten (1983) and Ezaz et al. (2005) as a guide for the expected number of chromosomes for final T2T assembly. A reference assembly was *generated* by choosing the best chromosome scaffolds from one of the two haplotype assemblies. The basis for selection was as follows for scaffolds >1Mbp in size. If the scaffold of haplotype 1 had both ends represented by telomeric sequence and the corresponding scaffold of haplotype 2 had only one end represented by telomeric sequence, then the scaffold for haplotype 1 was chosen for the reference assembly, and vice versa. If both the scaffolds for haplotype 1 and haplotype 2 contained telomeric sequence at both ends, then the scaffold with the fewest gaps was chosen for the reference assembly. If both T2T haplotypes had the same number of gaps, then the longest scaffold was selected for the reference assembly. If both haplotypes were equal in telomere presence, number of gaps and length, haplotype 1 sequence was chosen for the reference assembly. The Z and W specific scaffolds were added to the reference assembly. All scaffolds <1Mbp were drawn from haplotype 1 for the reference assembly.

#### Chromosome assignments

Bacterial Artificial Chromosome (BAC) clones were previously used for generating physical map for *Pogona vitticeps* (Young et al., 2013; Deakin et al., 2016). BAC end sequences (n=273) corresponding to 137 clones were downloaded from the NCBI GSS database. These sequences were aligned to the reference genome using *minimap2* (parameters: -x asm20 --secondary-no) to identify their locations in the reference genome. We also mapped the sex-linked sequence represented by 3,288 bp Clone C1 of Quinn et al. (2010) (Genbank accession EU938138) generated by walking out from a sex-linked 50 bp AFLP Pvi72W marker (Genbank accession ED982907) identified by Quinn et al. (2007) to confirm the assignment of a scaffold to the non-recombining region of the W chromosome (Scaffold 17).

#### Read depth and GC content calculations

PacBio HiFi (parameter: -x map-pb) and ONT (parameter: -x map-ont) sequence data were aligned to the scaffold assembly using *minimap2* (v2.17, Li 2018) Similarly, Illumina sequence data were aligned to the assembly using *bwa-mem2* (v2.2.1, Vasimuddin *et al*. 2019) using default parameters. Resulting alignment files were sorted and indexed for efficient access using *samtools* (v1.19, Danecek *et al*. 2021). Read depth in non-overlapping sliding windows of 10 Kbp was calculated using the *samtools bedcov* command. GC content in non-overlapping sliding windows of 10 Kbp was calculated using *calculateGC.py* script.

#### Telomere repeats

*Tandem Repeat Finder* (*TRF*) (v4.09.1, Benson 1999, parameters: 2 7 7 80 10 500 6 -l 10 -d -h) was used to detect all repeats up to 6 bp length. TRF output was processed using *processtrftelo.py* script to identify regions >600 bp that contained conserved vertebrate telomeric repeat motif (TTAGGG). These regions were labeled as potential telomeres.

#### Centromere annotations

Enrichment of satellite repeats, increased inter-chromosomal HiC contacts (Mokhtaridoost et al., 2024), and reduced recombination typically mark centromeric regions. To identify satellite repeats we followed the procedure described by Zhang et al. (2023) with some modifications. Briefly, we counted 101-mers occurring 20 times or more with k-mer counter *KMC* (v3.2.4, Kokot et al., 2017, parameters: k=101, ci=20, -cs=100000). Satellite Repeat Finder (*SRF*, Zhang et al. 2023, commit id e54ca8c) was used to identify putative satellite repeats using those k-mers. Identified repeat units were elongated up to 1000 bp if they were <1000 bp, and all-vs-all alignments were performed using *minimap2* to group repeats into classes based on their sequence similarity. The reference genome was aligned to the identified repeat units using *minimap2* (Li, 2018 Parameters: –c –N1000000 –f1000 –r100,100 <(srfutils.js enlong srf.fa)). Note that repeat units <200 bp were extended to 200 bp before alignments using the srfutils.js utility in *SRF*. Alignments were processed using *srfprocess.R* script to merge consecutive alignments to the same repeat unit separated by <10 bp. All regions >100 bp long and 10% of the repeat unit length were retained for further analysis. If a genomic region overlapped multiple repeat classes, the longer region with its repeat class was chosen as a set of putative satellite repeat region with corresponding repeat class.

HiC inter-chromosomal interactions were examined and quantified for their association with centromeres. HiC data were mapped against the reference genome using the *GEM mapper* (v3.6.1, Marco-Sola et al. 2012) from *TADbit* (v1.0.1, Serra et al. 2017). Reads were iteratively mapped using windows from 15 bp to 75 bp in 5 bp steps. Possible artifacts were then removed, including: “self-circle”, “dangling-end”, “error”, “extra dangling-end”, “too short”, “too large”, “duplicated” and “random breaks”. Binning and data normalization were conducted using an in-house script that imports the “HiC_data” module of *TADbit* to bin unique reads into a square matrix of 50 Kbp. A 500 Kbp matrix was created and subsequently processed with *HiCExplorer* (v3.7, Ramírez et al., 2018). Both 50 Kbp and 500 Kbp matrices were corrected with Iterative Correction and Eigenvector (ICE) decomposition and normalized to a total of 100,000,000 interaction counts by scaling the sum of all interactions within the matrix. Normalized matrices were then plotted at a 500 Kbp resolution using *HiCExplorer*. The normalized 50 Kbp matrix was transformed into a GInteraction table using *HiCExplore*r, which includes interaction values between all genomic bins. Inter-chromosomal interactions were log-transformed and normalized to obtain Z-score values for each chromosome and genomic bin, as previously described (Alvarez-Gonzalez et al. 2022; Bista et al. 2024). Z-score values were plotted with ggplot2 as points and the LOESS method (span=0.4, Cleveland, 1979) was used for best fit line.

For measuring heterozygosity changes across the genome, each haplotype sequence was aligned to the reference genome using *minimap2* (parameters: -x asm5 --cs –K 1000M). Resulting alignments were processed using *paftools.js call* to identify variant sites. Since one of the haplotype sequences is the reference sequence, all variable sites are considered as heterozygous sites. Heterozygous variant site counts in 50 Kbp windows were counted and plotted using *ggplot2*. LOESS smoothing (span=0.5) was applied for the best fit line.

#### Sex chromosome identification

The putative Z and W scaffolds will have half the read depth of the autosomal scaffolds in a ZW individual. Scaffolds >1 Mbp long were examined for median read depths in 10 Kbp windows. Sex specific Z and W scaffolds were identified by having approximately half the median read depth of autosomes and the PAR in the sequenced ZW individual (Figure 10). The PAR scaffold was identified by homology with known Z chromosome sequence.

#### HiC analysis for sex chromosome differences in contact maps

HiC reads were quality-trimmed using *Trimmomatic* v0.39 to remove adapter sequences and low-quality reads. The trimmed reads of HiC data and Illumina DNA sequence data were aligned to both genome haplotypes using *BWA-mem* (v0.7.17). PacBio data were aligned to both genome haplotypes using *minimap2* v2.28. Resulting BAM files were merged and coordinate-sorted using *SAMtools* v1.19.2. Variant calling for each haplotype was performed using Illumina BAM files with *FreeBayes* v1.3.8 to generate VCF files. These VCF files were normalized using *BCFtools* v1.14, then compressed using *bgzip* from *HTSlib* v1.20. Phasing of VCF files was then conducted using *WhatsHap* v2.3 to resolve haplotype-specific information across the dataset, using genome haplotypes, normalized VCF files and PacBio BAM files as inputs. Phased VCF files were then used to phase the mapped HiC reads.

#### Mitochondria genome assembly

PacBio HiFi and ONT sequences were aligned to a *Pogona vitticeps* reference (NCBI Accession: NC_006922, Amer and Kumasawa, 2005) using *minimap2* (parameters: --map-pb or --map-ont) to search for mitochondrial reads. Alignments were processed to identify reads <20 Kbp and aligned residues >5 Kbp. No PacBio HiFi reads were identified using this filter. ONT reads were assembled using *flye* (v2.9.3, parameters: --iterations 2, Kolmogorov et al., 2019) to generate mitochondrial genome sequence. The output assembly sequence was processed using *MitoHiFi* (v2.9.5, Uliano-Silva et al.,2023) to adjust the start coordinate and obtain annotations.

### Assembly evaluation

The assembly was evaluated against criteria established by the Earth Biogenomes Project (EBP, https://www.earthbiogenome.org/report-on-assembly-standards, version 6CQ40, Lawniczak et al., 2022) namely: percentage of collapsed sequence, percentage false expansions, k-mer completeness, complete single copy BUSCO genes, and average percentage of transcriptome data mappable to the genome assembly and contaminations (Figure 2).

#### K-mer completeness and per base error rate estimation

Illumina sequence data were trimmed for adapters and low-quality reads using *Trimmomatic* (v0.39, Bolger *et al*. 2014, parameters: ILLUMINACLIP:TruSeq3-PE.fa”:2:30:10:2:True LEADING:3 TRAILING:3 SLIDINGWINDOW:4:20 MINLEN:36). Resultant paired-end sequences were used to generate k-mer database using *meryl* (v1.4.1, Rhie *et al*. 2020). *Merqury* (v1.3, Rhie et al. 2020) was used with *meryl* k-mer database to evaluate assembly k-mer completeness and estimate per base error rate of pseudo-haplotype and individual haplotype assemblies.

#### False expansions and collapses

Putative false expansion and collapse metrics were calculated using the *Inspector* (v1.2, Chen et al., 2021, default parameters) and PacBio HiFi data.

#### Contamination check

Vector contamination was assessed using *VecScreen* defined parameters for *BLAST* (v2.14.1, Camacho et al., 2009, parameters: -task blastn -reward 1 -penalty -5 -gapopen 3 -gapextend 3 -dust yes -soft_masking true -evalue 700 -searchsp 1750000000000) and the *UniVec* database (accessed on 18^th^ June 2024).

#### Gene completeness evaluation

*BUSCO* (v5.4.7, Manni *et al*. 2021) was run using *sauropsida_odb10* library in offline mode to assess completeness metrics for conserved genes. BUSCO synteny plots were created with *ChromSyn* (v1.3.0, Edwards et al. 2022).

#### RNAseq mapping rate

RNAseq data from multiple tissues (Table S2) were aligned to the assembly using *subread-align* (v2.0.6, parameters: -n 150 Liao *et al*. 2013) to calculate percentage of mapped fragments for evaluating RNAseq mapping rate. We chose –n 150 to sample all possible seeds for alignments because of high heterozygosity observed for the species. We did not have RNAseq data for the focal individual used for the genome assembly.

### Annotation

#### Repeat annotation

*RepeatModeler* (v2.0.4, Smit et al. (2008-2015) parameters: -engine ncbi) was used to identify and classify repetitive DNA elements in the genome. Subsequently, *RepeatMasker* (v4.1.2-pl, Smit et al. (2013-2015) was used to annotate and soft-mask the genome assembly using the species-specific repeats library generated by *RepeatModeler* and families were labelled accordingly.

#### Ribosomal DNA

Assembled scaffolds were searched for ribosomal DNA units using *ribocop.py* which searches for consecutive alignments of 18S, 5.8S, and 28S to determine rDNA sequences.

#### De novo gene annotations

RNAseq data from multiple tissues (Table S2) were processed using *Trinity* (v2.12.0, Grabherr *et al*. 2011, parameters: --min_kmer_cov 3 --trimmomatic) to produce individual transcriptome assemblies. Parameters were chosen to remove low abundance and sequencing error k-mers. The assembled transcripts were aligned to the UniProt-SwissProt database (last accessed on 28-Feb-2024) using *diamond* (v2.1.9, Buchfink *et al*. 2021, parameters: blastx --max-target-seqs 1 --iterate --min-orf 30). Alignments were processed using *blastxtranslation*.*pl* script to obtain putative open reading frames and corresponding amino acid sequences. Transcripts containing both the start and the stop codons, with translated sequence length between 95% and 105% of the best hit to UniProt_SwissProt sequence, were selected as full-length transcripts.

Amino acid sequences of full-length transcripts were processed using *CD-HIT* (v4.8.1, Fu *et al*. 2012, parameters: -c 0.8 -aS 0.9 -g 1 -d 0 -n 3) to cluster similar sequences with 80% pairwise identity and where the shorter sequence of the pair aligned at least 90% of its length to the larger sequence. A representative transcript from each cluster was aligned to the repeat-masked genome using *minimap2* (v2.26, parameters: --splice:hq), and alignments were coordinate-sorted using *samtools*. Transcript alignments were converted to *gff3* format using *AGAT* (v1.4.0, Dainat, 2022, agat_convert_minimap2_bam2gff.pl) and parsed with *genometools* (v1.6.2, Gremme *et al*. 2013) to generate training gene models and hints for *Augustus* (v3.4.0, Stanke et al. 2008) with untranslated regions (UTRs). Similarly, transcripts containing both start and stop codons with translated sequence length outside of 95% and 105% of the best hit to UniProt_SwissProt sequence, were processed in the same way to generate additional hints. A total of 500 of these representative full-length transcripts were used in training for gene prediction to calculate species-specific parameters. During the gene prediction model training, parameters were optimized using all 500 training gene models with a subset of 200 used only for intermediate evaluations to improve run time efficiency. Gene prediction for the full dataset used 20 Mbp chunks with 2 Mbp overlaps to improve run time efficiency.

An issue was identified where the predicted *Amh* gene on the Z-specific scaffold (scaffold 18) was fused with neighboring genes. To resolve this, gene prediction was rerun on the Z scaffold with manually modified hints. Specifically, the weighting of UTR hints intersecting with the two predicted introns flanking the *Amh* coding sequence was increased to down weight intronic predictions by *Augustus* in that region. The updated Z scaffold gene predictions were then concatenated with the original gene predictions.

Predicted genes were aligned against Uniprot_Swissprot database for functional annotation using best-hit approach and *diamond*. Unaligned genes were subsequently aligned against Uniprot_TrEMBL database for functional annotation.

## Results and Discussion

### DNA sequence data quantity and quality

PacBio HiFi sequencing yielded 70.6 Gb with a mean read length of 14,980 bp (Table 1) and mean quality value >Q30 of all reads. The ONT sequencing yielded 105.6 Gb of reads with an N50 value of ~37 Kbp and 48.1% reads with mean quality value >Q20 (Table 2). The distributions of quality scores and read lengths for the long-read sequencing align with known characteristics of the ONT and PacBio platforms (Figure S1). K-mer frequency histograms of Illumina, ONT and PacBio HiFi sequence data for k=17, k=21 and k=25 show two distinct peaks (Figure 3) confirming the diploid status of this species. The peak for heterozygous k-mers was smaller for k=17 compared to the homozygous k-mer peak. In contrast, the heterozygous k-mer peak was higher for k=25 compared to the homozygous k-mer peak, suggestive of high heterozygosity at a small genomic distance. Genome size was estimated to be 1.81 Gb using the formulae of Georges *et al*. (2015) and Illumina sequence data, with a k-mer length of 17 bp, homozygous peak of 45.5 (Figure 3) and the mean read length of 134.3 bp. However, the PacBio estimate of genome size of 1.74 Gb agrees more closely with the previous estimate using earlier Illumina reads (Georges et al. 2015) and the estimate from flow cell cytometry of 1.77 Gb (Georges et al. 2015). The reason for the discrepancy between the current and former estimates of genome size from Illumina data is unclear, but may have arisen because the current Illumina data was not filtered for error reads in the same way.

**Table 1.** Summary metrics for sequence data and assembly for the bearded dragon *Pogona vitticeps*.

| Sequencing Platform | Number of Reads | Mean Read Length (bp) | Total Bases | Est. Genome size (Gbp) | Read Depth |
| --- | --- | --- | --- | --- | --- |
| Illumina PE DNA | 694,401,150 | 134 | 93,255,202,850 | 1.81 | 52.6x |
| PacBio HiFi Sequel II | 4,714,654 | 14,980 | 70,625,888,904 | 1.74 | 40.4x |
| ONT R9.4.1 | 7,118,515 | 14,830 | 105,566,275,033 | 2.7 | 60.5x |
| Arima Genomics HiC | 590,697,330 | 151 | 89,195,296,830 | -- | -- |

**Table 2.** Summary metrics for the genome assembly of the bearded dragon *Pogona vitticeps*. The pseudo-haplotype is a combination of haplotypes 1 and 2 (sensu *hifiasm*); the Reference Assembly was constructed by selecting the best scaffolds from each of haplotypes 1 and 2.

| Metric | Haplotype 1 | Haplotype 2 | Pseudo-haplotype | Reference Assembly |
| --- | --- | --- | --- | --- |
| Assembly length | 1,752,200,003 | 1,747,167,247 | 1,747,460,405 | 1,752,814,424 |
| No. of scaffolds/contigs | 89 | 51 | 71 | 89 |
| GC Content | 42.2 | 42.2 | 42.2 | 42.2 |
| No. of gaps > 100 bp | 31 | 28 | 15 | 26 |
| Mean sequence length | 19,687,640 | 34,258,181 | 24,612,118 | 19,694,544 |
| Median sequence length | 77,000 | 152,108 | 85,021 | 77,000 |
| Longest sequence | 359,918,989 | 358,276,425 | 359,349,958 | 358,276,425 |
| Shortest sequence | 4,000 | 18,330 | 4,000 | 4,000 |
| N50 | 265,980,915 | 266,210,064 | 266,029,613 | 266,210,064 |
| N90 | 28,115,431 | 28,121,876 | 28,118,385 | 28,115,431 |
| L50 | 3 | 3 | 3 | 3 |
| L90 | 9 | 9 | 9 | 9 |

**Figure 3.**
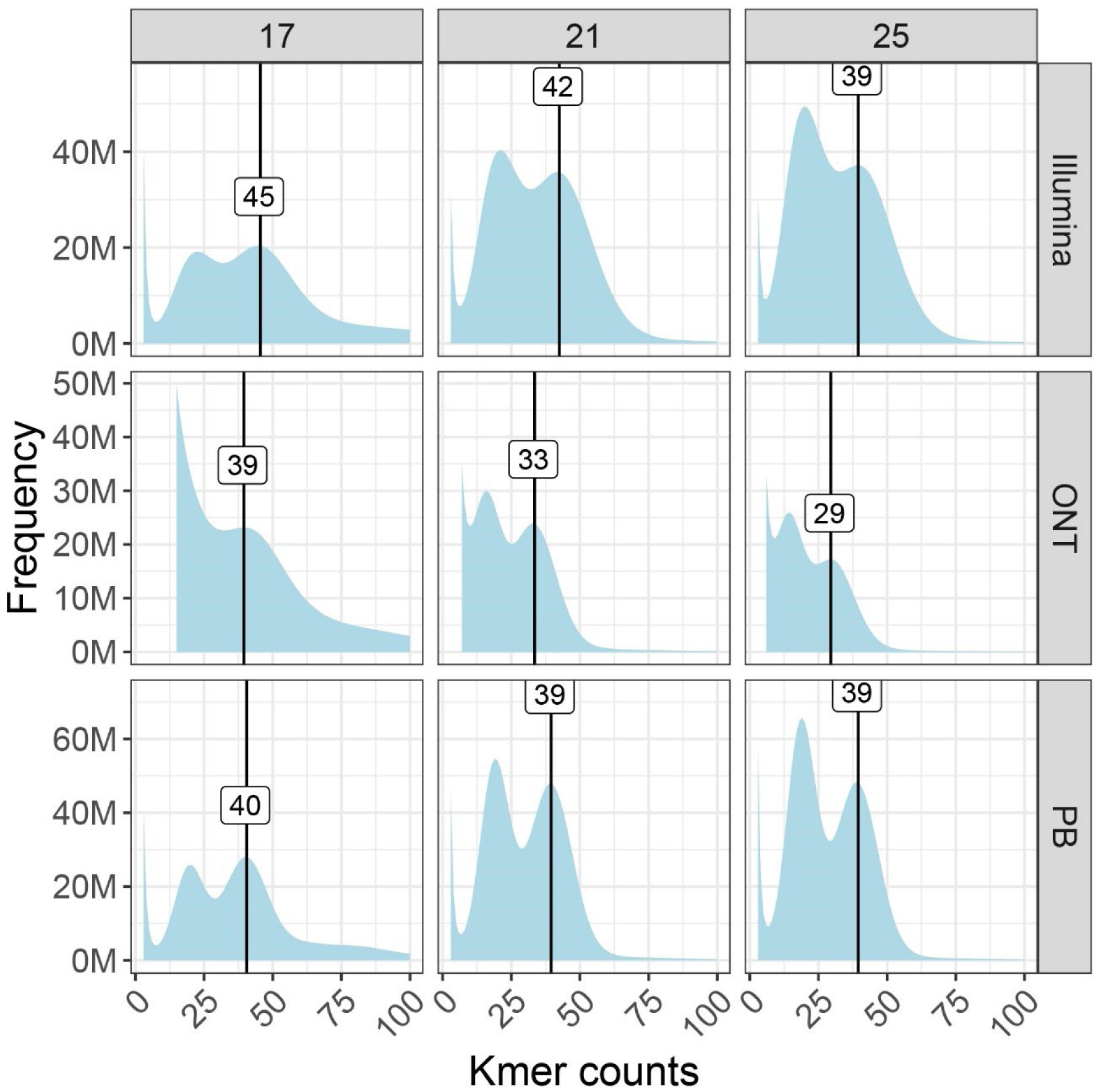
Distribution of k-mer counts frequency using sequences from Illumina, Oxford Nanopore Technologies (ONT), and PacBio (PB) platforms for the bearded dragon *Pogona vitticeps*. Heterozygosity is high as indicated by dual peaks in each graph, and the height of the heterozygous peak increases with the length of the k-mer. This confirms diploidy.

Read depth, obtained by dividing the total DNA sequence data from each platform by the assembly size, was consistent (Table 1) with the median read depths of 60.6*x* for ONT, 40.5*x* PacBio HiFi and 52.7*x* Illumina platforms calculated for 10 Kbp non-overlapping sliding windows of the assembly.

### Assembly

*Hifiasm* produced three assemblies: one for each haplotype and a pseudo-haplotype of high quality as evidenced by assembly metrics (Table 2). The haplotype assemblies were subject to further scaffolding and mis-join error correction using the HiC data to improve assembly contiguity (Figure S2). Minimal manual curation was required as wrongly joined scaffolds were corrected by YaHS (Figure S2). The reference assembly for the central bearded dragon had a total length of 1,752,814,424 bp assembled into 89 scaffolds, with 26 gaps each marked by 100 Ns. This compares well with other published squamate genome assemblies.

The central bearded dragon reference genome (PviZW2.1) is contiguous with a scaffold N50 value of 266.2 Mbp and a N90 value of 28.1 Mbp with the largest scaffold of 358.3 Mbp (Table 2). L50 and L90 values were 3 and 9 respectively, typical of species with microchromosomes, where most of the genome is present in large macrochromosomes.

All 15 major scaffolds in the assembly (corresponding to autosome number in the karyotype of the bearded dragon) had well defined telomeres at each end (Figure 4). Telomeres were comprised of the vertebrate telomeric motif TTAGGG and ranged in size from 2,430 bp (405 copies of the repeat motif) to 42,098 bp (7,151 repeat copies). The telomeric regions were typically characterized by an expected rise in GC content (Figure 4) and a significant rise in inter-chromosomal contact (Figure 5; Figure S3), mirroring patterns previously described in turtles (Bista et al. 2024).

**Figure 4.**
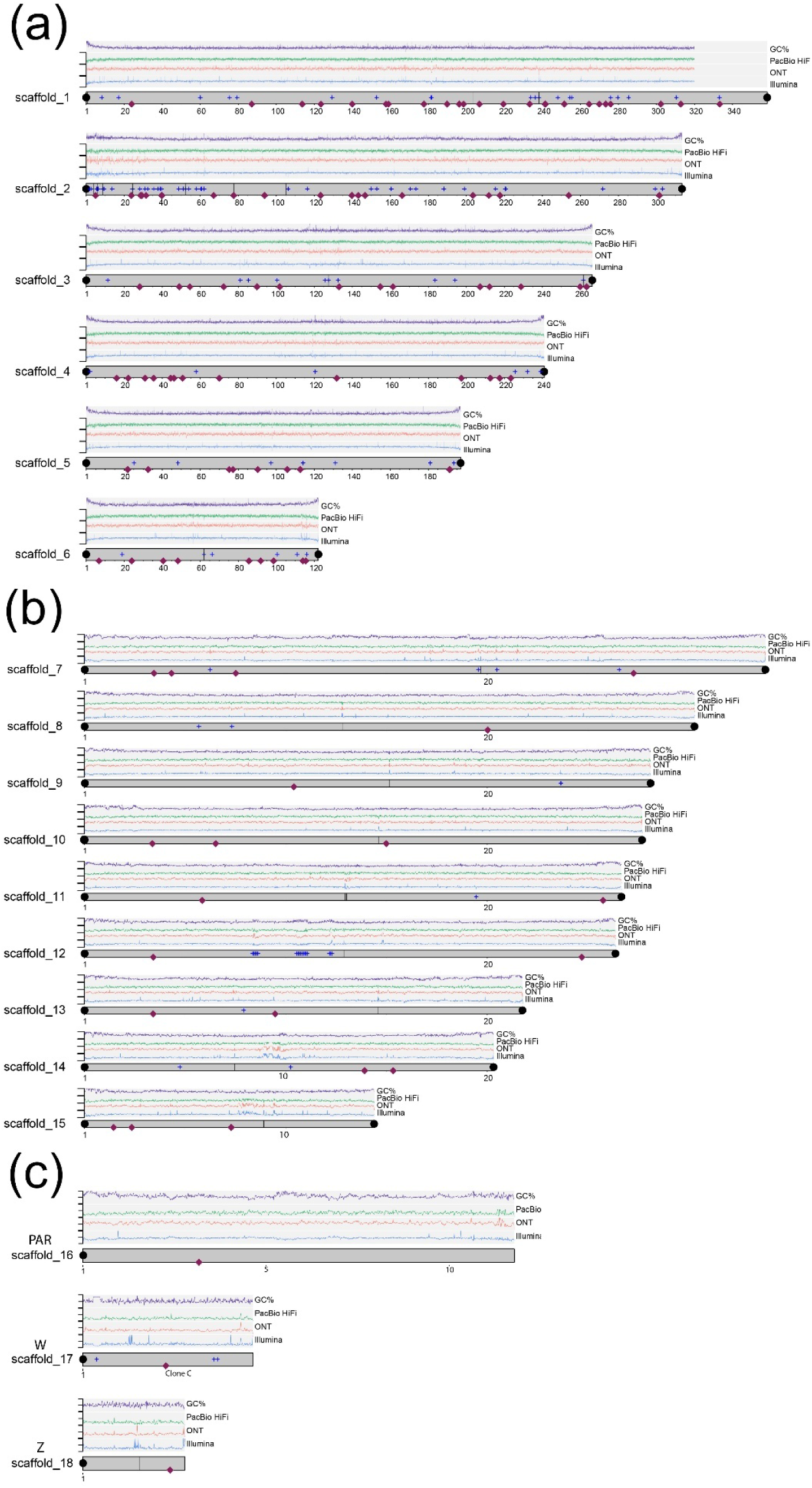
A plot of the 18 longest scaffolds (corresponding to the number of chromosomes of the bearded dragon *Pogona vitticeps*. Four traces are shown on each chromosome. The top trace (purple, range 30-60%) represents GC content, the next trace (green, range 0-50*x*) represents PacBio HiFi read depth, the next trace (red, range 0-100*x*) represents ONT read depth, and the fourth trace (blue, range 0-100*x*) represents Illumina read depth. Note that there is no indication in any of these traces of centromeric position in contrast to *Bassiana* (Hanrahan et al., 2025). Telomeres are shown as black dots; satellite repeats are indicated by the blue plus symbols (+); gaps by vertical black lines. The red diamonds show the location of BAC anchors (Young et al., 2013; Deakin et al., 2016, Table S7). Locations of the putative centromeres are shown in Figure 5. (a) Macrochromosomes; (b) Microchromosomes (c) both the Z and W specific regions were assembled into single scaffolds, with the PAR assembled into a single scaffold in both haplotypes. Refer to Supplementary Materials for a high-resolution version of this figure.

**Figure 5.**
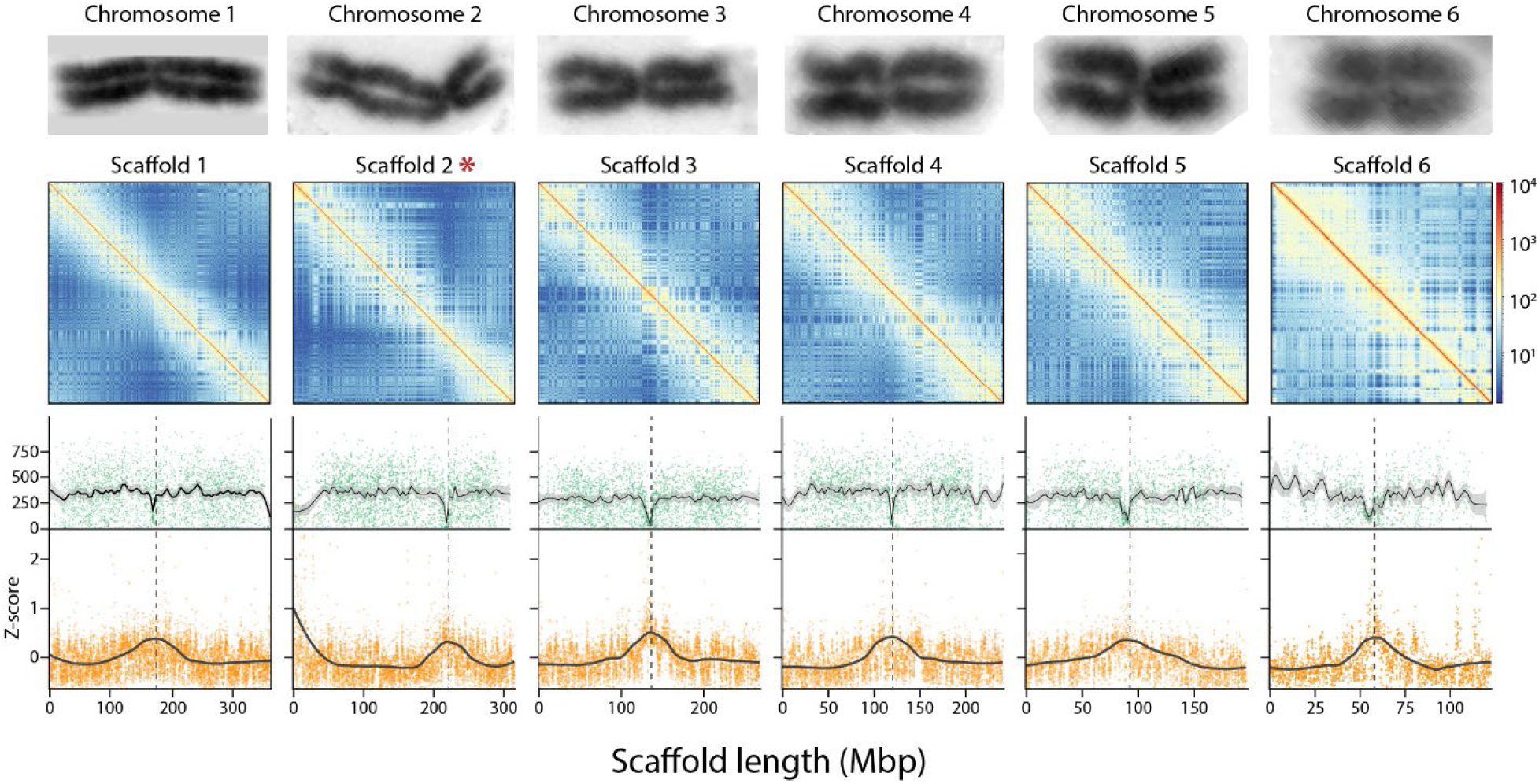
Identification of putative centromeres for the six macrochromosomes. The upper row of panels gives chromosome-specific Hi-C heatmaps showing intra-chromosomal interactions. The second row of panels shows the count of heterozygous sites per 50 Kbp window (green dots) with lines of best fit and 95% confidence interval (grey shading). The lower row of panels are the Z-scores for inter-chromosomal HiC interactions along chromosome length (Mbp) with smoothed lines of best fit. Each dot in the lower panels represents the Z-score interaction value of a different 50 Kbp bin. Chromosomes images are taken from Ezaz et al. (2005) and are not to scale. They are to illustrate the correspondence between the karyotype centromere and the putative position of the centromere (dashed lines) inferred from the dip in heterozygosity and the peak in inter-chromosomal contact. Scaffold 2 marked (*) is inverted with respect to the published karyotype. Refer to Figure S3 for similar plots for the microchromosomes.

Initially, we did not detect telomere repeat sequence on 5’ end of the Scaffold 10 using a stringent threshold of 600 bp for telomeric region. However, manual examination revealed 32 repeats of telomeric sequence from position 1-214 on Scaffold 10 verifying that it had telomeres at both ends. The missing telomere for Scaffold 16 is expected because it is the pseudo-autosomal region of the sex chromosomes. The putative sex chromosome scaffolds 17 = W and 18 = Z also each possessed only one terminal telomeric sequence. This is consistent with T2T assembly for the sex chromosomes once the PAR and the non-recombining regions of Z and W are combined.

Typical centromeric satellite repeats units were not evident in the repeat structure, read depth profiles or GC content profiles (Figure 4) as they were for *Bassiana duperreyi* (Hanranan et al. 2025). Putative centromeric regions were evident for the macrochromosomes as an increase in the levels of inter-chromosomal contact in the HiC data and as a drop in heterozygosity (Figure 5 and Figure S3).

Of 137 BAC clones (Young et al., 2013; Deakin et al., 2016), 5 with single sequences did not align, 2 had inter-chromosomal mappings, 14 had discrepant mappings for macrochromosomes and 2 had end sequences that were too far apart to be considered valid. This left 114 clones (83.2%) with reliable mappings. This physical mapping validated the assignment of assembly scaffolds 1-6 to the macrochromosomes 1-6 of the genome (Figure 4a). The assignment of scaffolds 7-15 to the microchromosomes (Figure 4b) albeit with altered order (Figure 7), scaffold 16 to the PAR of the sex chromosomes (Figure 4c), and scaffold 18 as the nonrecombining region of the Z chromosome (Figure 4c). Mapping of the W-linked sequence Clone C1 (3,288 bp, Quinn et al., 2010) confirmed the identity of scaffold 17 as the non-recombining region of the W chromosome (Figure 4c).

**Figure 6.**
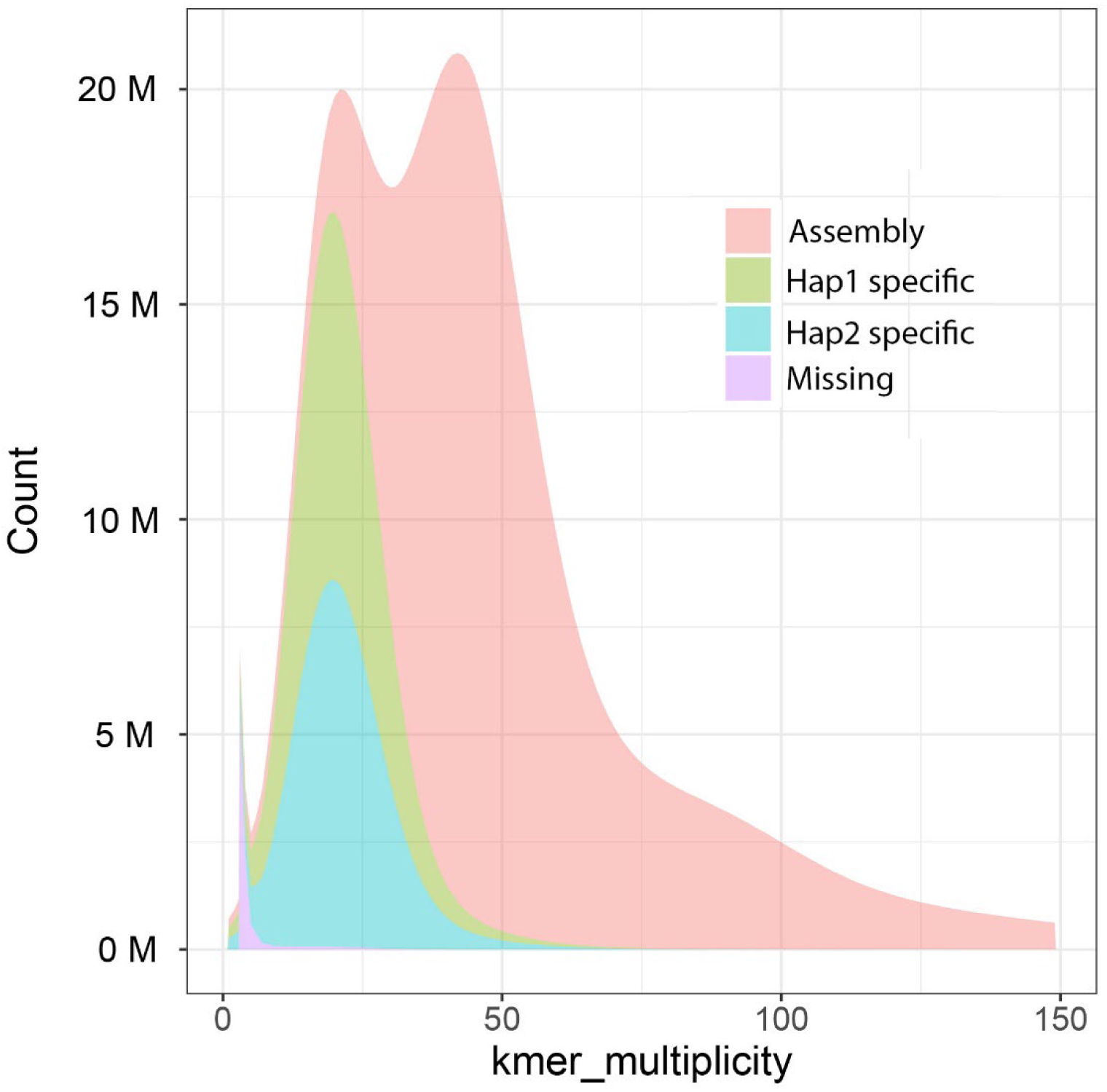
Distribution of Illumina k-mers (k = 17) in the genome assembly of the bearded dragon *Pogona vitticeps* (Table S6). K-mer counts are shown on the x-axis and the frequency of occurrence of those counts on the y-axis. Those scored as missing are found in reads only.

**Figure 7.**
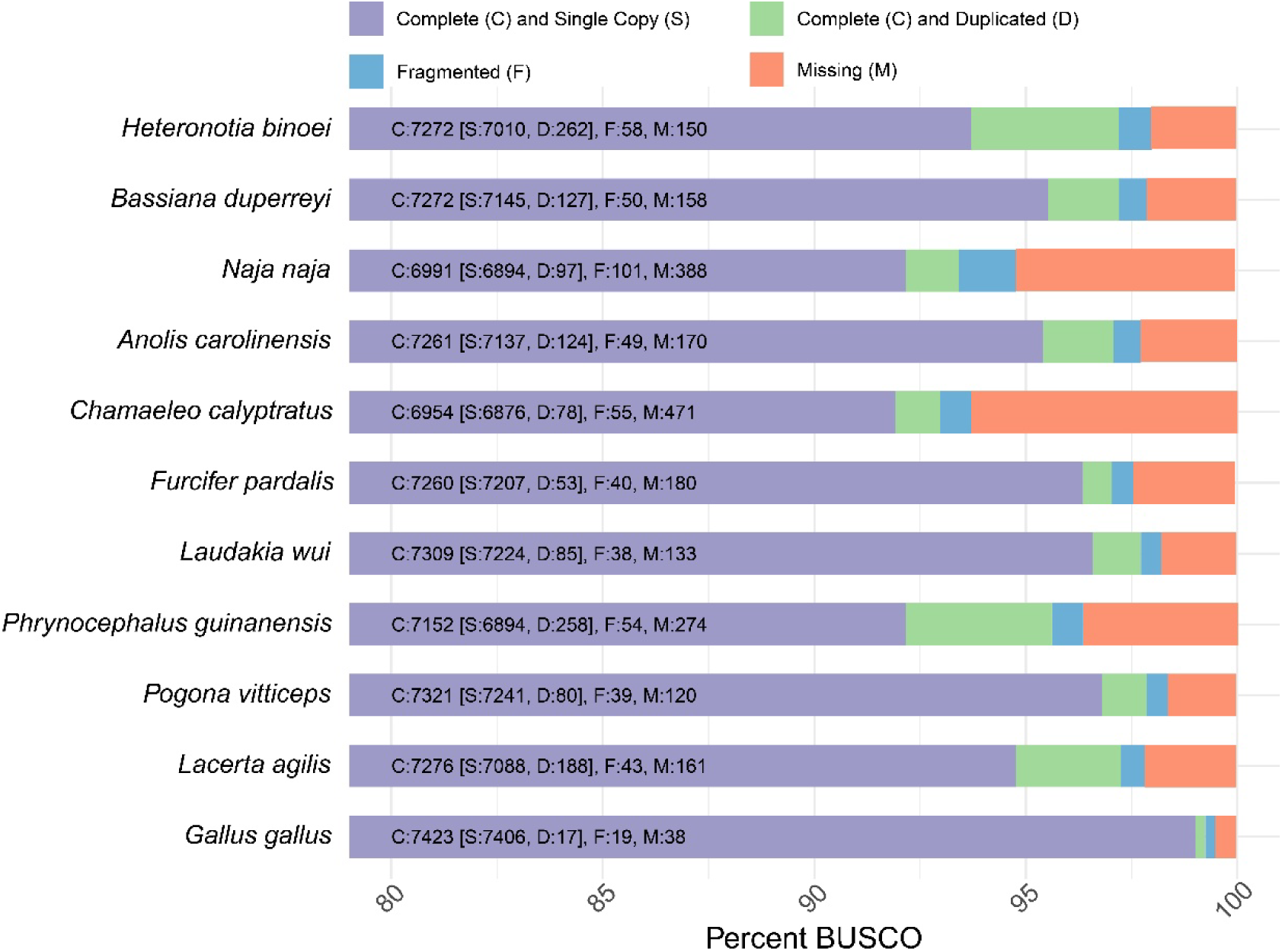
A visual representation of how complete the gene content is for each listed species genome, including *Pogona vitticeps*, based on Benchmarking Universal Single-Copy Orthologs (BUSCO, n=7480).

### Assembly evaluation

The percent collapsed sequence in the assembly was exceptionally low at 0.003% (492,971 bp, 54-13,643 bp, n=255) as was the percentage of false expansions at 0.03% (49,447 bp, 52-5,133 bp, n=69); two of the indicators of genome assembly quality identified by the Earth Biogenome Project (Lawniczak et al., 2022).

Completeness of the assembly was estimated to be 99.82% for both haplotype assemblies combined and the per base assembly quality estimate exceeded Q40 at 48.36 (1 error in 146 Kbp). High heterozygosity in the k-mer profiles (Figure 3) affects assembly completeness metrics measured by *Merqury*. Individual haplotype assemblies were 85.5% complete, which is expected of animals with high heterozygosity (in our case, 1.98%). This shows that assembly completeness metrics for a single haplotype assembly measured using k-mers can be understated for species with high heterozygosity.

Analyses using the Benchmarking Universal Single-Copy Orthologs (BUSCO) gene set for Sauropsids reveals 7,321 genes as complete (97.9%), with a minimal proportion duplicated (D: 1.1%), indicating a robust genomic structure with minimal redundancy (Figure 7). The central bearded dragon genome also had a low proportion of fragmented (F: 0.5%) and missing (M: 1.6%) orthologs. These results positioned central bearded dragon favorably in terms of genome completeness and integrity, on par with other squamates, and highlights its potential as a reference for further genomic and evolutionary studies within this phylogenetic group. In our comparison set, only chicken (*Gallus gallus*) has better BUSCO statistics than the bearded dragon. RNAseq data mappability was on average 93.5% and 18 of 22 samples had more than 90% of fragments mapped to the genome (Table S2). Note that sensitivity settings for alignments had to be increased for mapping RNAseq data given high hetergozygosity observed for this species (1.98%).

### Chromosome Assembly

The bearded dragon has 2n=32 chromosomes with six pairs of macrochromosomes and ten pairs of microchromosomes including the sex chromosomes. The distinction between macro and microchromosomes typically relies on a bimodal distribution of size, however other characteristics such as GC content provide additional evidence for this classification (Waters et al. 2021; Bista et al., 2024) (Figure 8). The median GC content of 10 Kbp windows for the six largest scaffolds (representing macrochromosomes) ranged between 40.7% and 41.8%. In contrast, the remaining 12 scaffolds ordered by decreasing length had a median GC content of between 42.6% and 47.6% characteristic of microchromosomes in other squamates.

**Figure 8.**
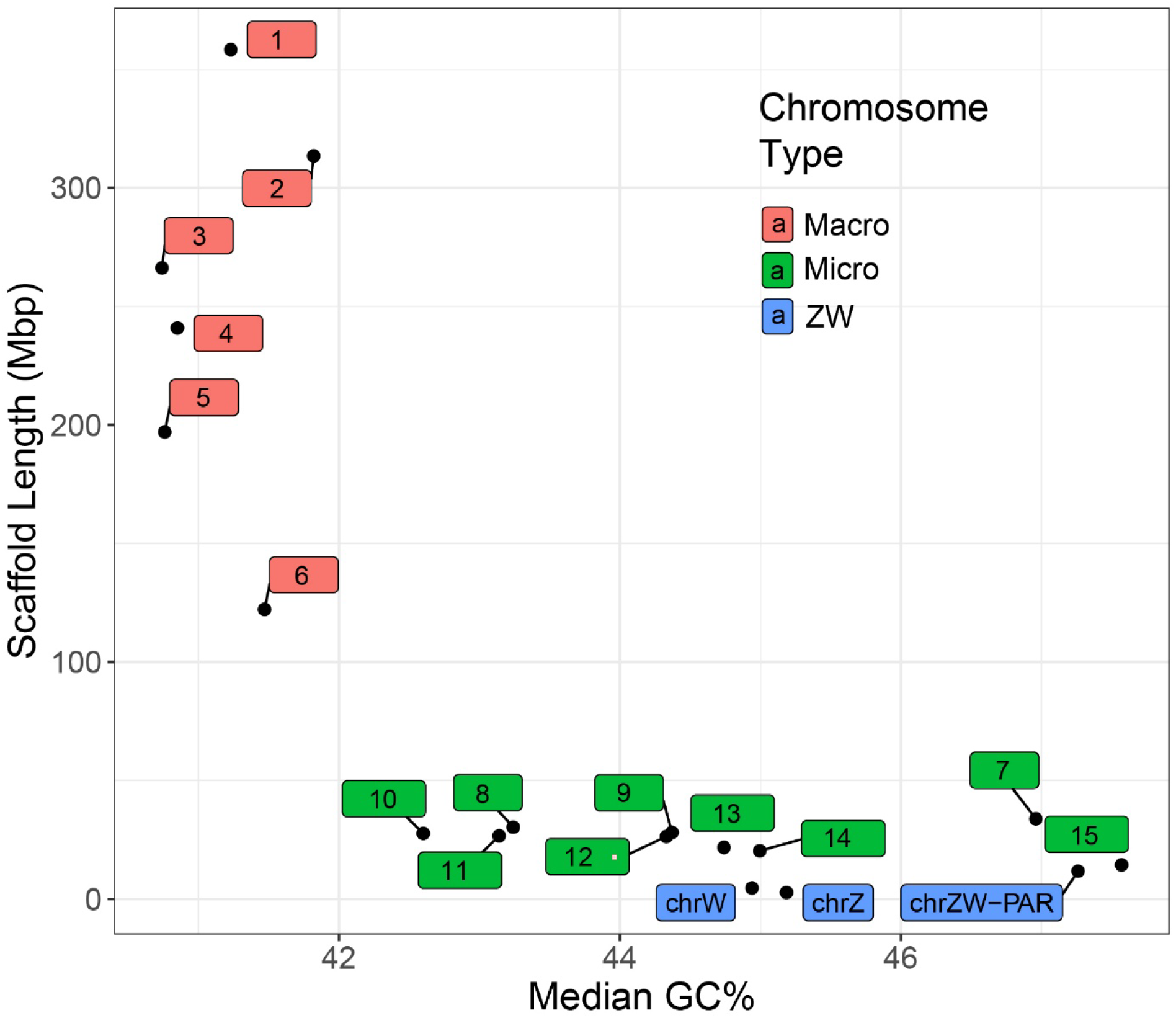
A plot of assembly scaffolds defined by scaffold length vs median GC content in 10 Kbp windows. Microchromosomes are characterised by higher GC content than macrochromosomes. Median GC content in 10 Kbp windows of scaffolds vs length of scaffolds representing macrochromosomes (scaffolds 1-6, red), the sex chromosomes (blue, the PAR and nonrecombining regions of the Z and W) and the other microchomosomes (green, scaffolds 7-15). Scaffold numbers 1-6 correspond to the macrochromosome numbers of Deakin et al. (2016) for scaffolds. Scaffold numbers 7-15 translate to the microchromosome numbers of Deakin et al. as per Table S6.

Unlike mammals, reptiles (including most birds) show a high level of chromosomal homology across species (Waters et al. 2021; Bista et al. 2024). Figure 9 shows synteny conservation between bearded dragon, representative squamate species and chicken. Apart from a handful of intrachromosomal rearrangements, the major scaffolds of bearded dragon and other squamates corresponded well, including the pseudoautosomal region (PAR) of the sex microchromosomes (scaffold 16) within the Agamidae. When compared with other genomes in the analysis, the bearded dragon genome showed a high degree of evolutionary conservation with respect to both chromosomal arrangement and gene order (Figure 9).

**Figure 9.**
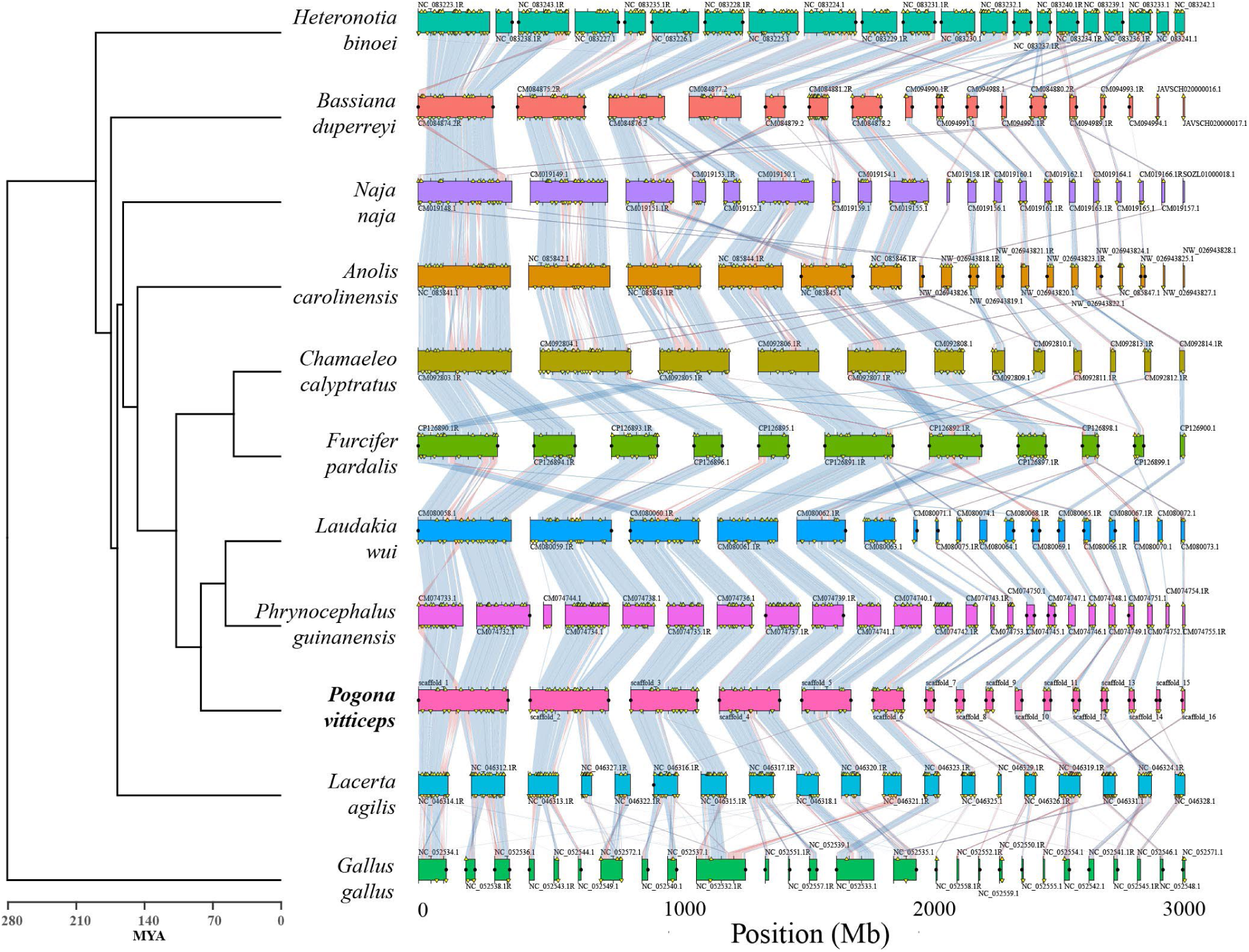
Synteny conservation of BUSCO homologs for the bearded dragon *Pogona vitticeps* and squamates with chromosome level assemblies including representative skink, iguanid, snake and gecko lineages and chicken. Synteny blocks corresponding to each species are aligned horizontally, highlighting conserved chromosomal segments across the genomes. The syntenic blocks are connected by ribbons that represent homologous regions shared between species, with the varying colours denoting segments of inverted gene order. Duplicated BUSCO genes are marked with yellow triangles. Predicted telomeres are marked with black circles.

The Z and W specific sex chromosome scaffolds were identified as 18 and 17, respectively. These represent the non-recombining region of the sex chromosomes. They were not assembled to the PAR in either haplotype. The Z specific scaffold was 2.78 Mbp and W specific scaffold was 4.64 Mbp. In the sequenced ZW female, read depth for both scaffolds were identified based on the median read depth in 10 Kbp sliding windows. As expected, read depth was approximately half that of the autosomes and the PAR scaffold (Figure 10a). The first half of the Z and W scaffolds share good homology (Figure 10b). On the second half of the W scaffold there appears to have been duplication and expansion that increased its size relative to the Z. The PAR scaffold (scaffold 16 reference, 11.77 Mbp) was identified by homology to known Z sequences from Pvi1.0.

**Figure 10.**
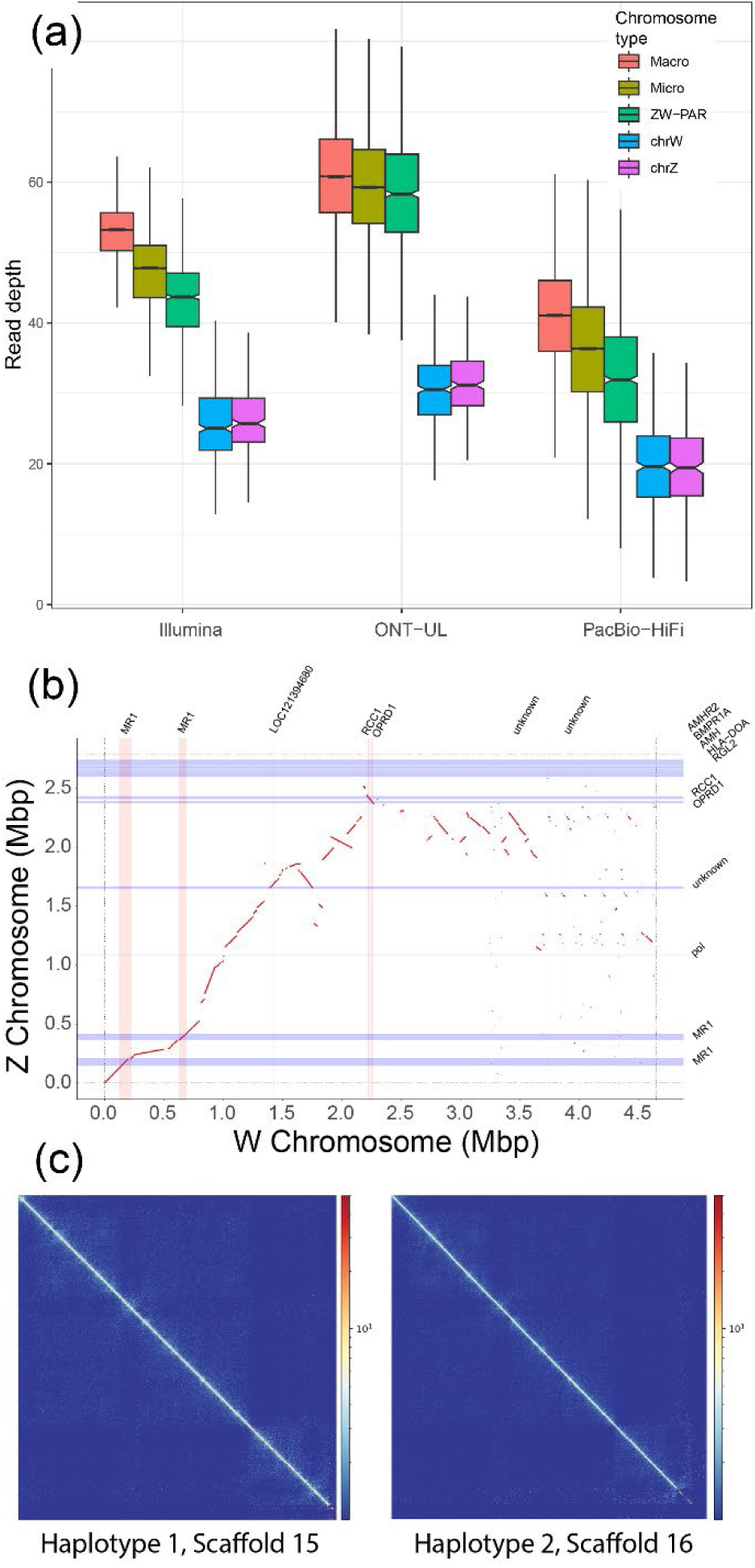
Sex chromosome analysis. a) For each sequencing technology, boxplots of read depth in 10 Kbp windows of macrochromosomes, microchromosomes, the PAR, and Z and W specific scaffolds. Boxes represent the middle 50% of the data, notches represent 95% confidence intervals of the medians (central horizontal black bar), whiskers are +/- 1.5 the interquartile range, outliers not plotted. b) Alignment of the Z specific (y-axis) and W specific (x-axis) scaffolds. Red lines represent homologies. Blue horizonal bars are genes annotated on the Z scaffold (gene names given on the y-axis), pink vertical bars are genes annotated on the W scaffold (gene names given on the x-axis). c) Phase HiC contact maps of the PAR in the two different haplotypes. Note that it is unknown which is the Z PAR and which is the W PAR.

The W specific scaffold had seven annotated genes, with none presenting as an obvious sex determining candidate. The Z specific scaffold also had seven annotated genes, four of which were ZW shared (Figure 10b). Notably, copies of both *Amh* and its receptor (*AmhR2*) were located on the Z, presumably duplicated from the autosomal homologues which remain present on scaffolds 7 and 2 respectively. Both genes are central to the sex determining pathway in other vertebrates, so present as strong sex determining candidates that would presumably function in a dosage dependent manner.

HiC reads were phased to the PAR scaffolds to determine if there was different 3D structure of the Z and W scaffolds (see Zhang et al., 2022). Surprisingly, despite clear cytological differences between the Z and W in cultured fibroblasts (Ezaz et al., 2005), there was little difference between HiC contact maps for each haplotype (Figure 10c). The discrepancy between cytogenetic (fibroblasts) and HiC data (blood) with respect to the Z and W structure, likely arise from the different cell types examined. Alternatively, the cytogenetic data only captures cells in metaphase when three-dimensional genome structure differences might be at their most pronounced.

### Annotation

#### General Repeat Annotation

An estimated 45.6% (798 Mbp) of the bearded dragon genome was composed of repetitive sequences, including interspersed repeats, small RNAs and simple and low complexity tandem repeats (Table 9). Retroelements (SINEs and LINEs) were the most common repetitive element (17.3%). DNA transposons were the second most common repetitive element (5.1%) and are dominated by Tc-Mar and hAT elements (Table S8). CR1, BovB and L2 elements were the dominant long interspersed elements (81% of LINE elements; 13.4% of the genome), which is consistent with other squamate genomes (Pasquesi et al. 2018). A total of 37% of all repeat content was unclassified and did not correspond to any element in the RepeatModeler libraries. Refer to Figure S4 for the size distribution of these unclassified repeats. The number of elements masked and their relative abundances are presented in the supplementary material (Table S8).

**Table 9.**
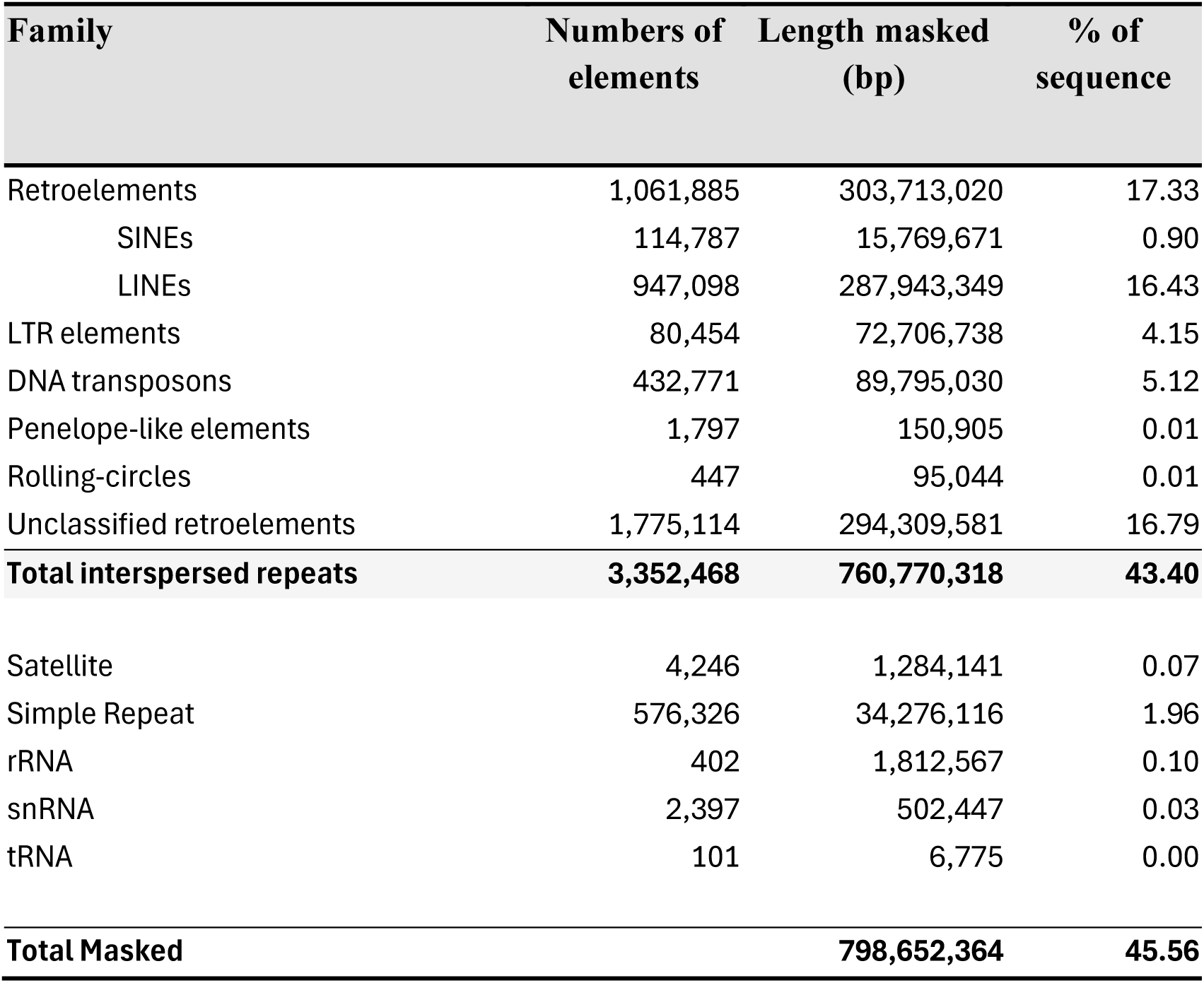
Condensed summary of the copy number and percentage of the bearded dragon (*Pogona vitticeps*) genome covered by repeat elements. Refer to. Table S8 **for full breakdown.**

| Family | Numbers of elements | Length masked (bp) | % of sequence |
| --- | --- | --- | --- |
| Retroelements | 1,061,885 | 303,713,020 | 17.33 |
| SINEs | 114,787 | 15,769,671 | 0.90 |
| LINEs | 947,098 | 287,943,349 | 16.43 |
| LTR elements | 80,454 | 72,706,738 | 4.15 |
| DNA transposons | 432,771 | 89,795,030 | 5.12 |
| Penelope-like elements | 1,797 | 150,905 | 0.01 |
| Rolling-circles | 447 | 95,044 | 0.01 |
| Unclassified retroelements | 1,775,114 | 294,309,581 | 16.79 |
| **Total interspersed repeats** | **3,352,468** | **760,770,318** | **43.40** |
| Satellite | 4,246 | 1,284,141 | 0.07 |
| Simple Repeat | 576,326 | 34,276,116 | 1.96 |
| rRNA | 402 | 1,812,567 | 0.10 |
| snRNA | 2,397 | 502,447 | 0.03 |
| tRNA | 101 | 6,775 | 0.00 |
| **Total Masked** |  | **798,652,364** | **45.56** |

### Satellite repeats

We undertook a more detailed analysis in an attempt to identify centromeric satellite repeats and centromeric regions as we did for the genome assembly of the skink *Bassiana duperreyi* (Hanrahan et al., 2025). The 67 satellite repeat units identified in the KMC/SRF analysis had lengths between 5 bp and 9,460 bp. These collapsed into 45 distinct classes based on sequence similarity (Table S9).

One repeat class (srfclass-16) corresponded to the telomeric microsatellite repeats (TTAGGG). A second class (srfclass-18) with a large repeat unit of 9,460 bp corresponded to ribosomal DNA sequence, dealt with in more detail later. A class of interspersed repeat (srfclass-11), possibly LINE elements, was comprised of 5,695 bp in unit length. A class of repeat (srfclass-30) was present as 84 copies on Scaffold 4 (119,114,420-119,423,712); all other copies were interspersed across the genome. A telomeric repeat was embedded in this larger repeat. A fifth class (srfclass-21) comprised repeat units of 2,190 bp on Scaffold 1 (117 copies, 167,719,827-167,977,892) and was somewhat enigmatic. These units were tandemly organized as 1 to 43 repeats, occasionally with a small intervening sequence. This repetitive sequence was found also on other scaffolds as interspersed units comprising a 1,406 bp motif and a 606 bp motif separated by 500 bp intervening sequence. A sixth class (srfclass-38) comprised repeat units of 877 bp, found only on scaffold 1 (238 copies, 54,463,716 – 254,497,379). A seventh class (srfclass-15) comprised repeat units of 398 bp, each as a composite of a 68 bp subunit, on scaffold 1 (181407117-181830275, *ca* 1063 copies). These repeat units align with elements on Scaffolds 3, 4 and 5, but with abbreviated subunits (e.g. 56 bp on Scaffold 3; 64 bp on Scaffold 4). An eighth repeat class (srfclass-4) comprised a 98 bp motif occurred on the W chromosome scaffold 17 (1,961 copies, 281,785-473,916) found as 109-450 bp alignments on other scaffolds. Of the 45 repeat classes, only one (srfclass-5, 151 bp) showed potential as a centromeric repeat unit. However, this repeat class was not distributed as a single consolidated cluster on each chromosome, as would be expected of centromeric repeat units.

We were thus unable to definitively identify centromeric repeat units in the bearded dragon to confirm the presence of only one per chromosomal scaffold as we were able to do in the skink *Bassiana duperreyi* (Hanrahan et al., 2025). We were however able to confirm the likely presence of one centromere per scaffold as expected if the scaffolds correspond to chromosomes using plots of heterozygosity and an index of inter-chromosomal contact rates against position on the scaffold (Figure 5). A dip in heterozygosity corresponded with a peak in HiC inter-chromosomal contact rate which together corresponded well with the position of the centromere taken from metaphase chromosomal spreads (Ezaz et al., 2005).

### Gene Annotation

We assembled transcriptomes from 22 samples (Table S2). Genome annotation using *Augustus* predicted 17,237 genes and transcripts, of which 16,799 had a match to a Uniprot_Swissprot or Uniprot_TrEMBL protein sequence, and 16,483 were assigned a gene name. The quality of the annotation was further validated using RNAseq data from 22 samples, with an average 54.4% (ranging from 22.8% to 76.4%) of aligned reads assigned to annotated exons, indicating a reasonable level of correspondence between the predicted gene models and the observed transcriptomes.

### Mitochondrial Genome

The bearded dragon mitochondrial genome assembly was 16,731 bp in size with 37 intact genes without frameshift mutations. It consisted of 22 tRNAs, 13 protein coding genes, 2 ribosomal RNA genes and the control region (Figure S5), so was typical of the vertebrate mitochondrial genome. Base composition was A = 33.0%, C = 29.8%, G = 13.1% and T = 24.0%.

We note that mitochondrial sequence was absent in the HiFi data presumably because it was eliminated during the size selection step. As the assembly software uses PacBio HiFi for the core assembly, these mitochondrial sequences, although present in the ONT data, were not recovered during the combined assembly process. We also note a drop in the read depth for the PacBio HiFi and Illumina data for exceptionally small microchromosomes (Figure 10a) that is not observed for ONT data. This suggests a systematic bias in the data from sequence-by-synthesis platforms for small elements and high GC content sequences.

### Ribosomal DNA

The rDNA unit length in the bearded dragon is approximately 9.5 Kbp, with a total of 1.75 Mbp of sequence across 24 scaffolds containing rDNA sequences. The rDNA sequence was found on chromosome 2 scaffold as expected (Young et al., 2013) near the sub-telomeric region of 2q. There were 23 additional short scaffolds comprised entirely of rDNA arrays as well indicating poor quality assembly of rDNA array. The first and second internal transcribed spacers (533 bp and 344 bp respectively) and intergenic spacer (2.7 Kbp) are relatively small, compared to mammals (McDonald et al., 2024).

## Conclusion

Here we present a high-quality genome assembly of the central bearded dragon, *Pogona vitticeps* (Ahl, 1926). The quality of the genome assembly and annotation compares well with other chromosome-length assemblies and is among the best for any species of Agamidae. We have chromosome length scaffolds, telomere-to-telomere.

The non-recombining regions of the Z and W chromosomes were each assembled as a single scaffold. The PAR was assembled as a single scaffold in both haplotypes. The sex chromosomes scaffolds and PAR scaffold each lacked one telomere, but this is likely resolved when they are combined to form Z and W scaffolds including both the PAR and non-recombining regions. The identification of *Amh* and *Amhr2* on the Z specific scaffold (but not the W) has them as strong candidates for the sex determining gene(s) in this species. Gene *Nr5a1,* encoding transcription factor SF1, was previously identified as a candidate sex determining gene because it resided on the sex chromosomes and because of its differential transcript isoform composition (Zhang et al., 2022); it is confirmed as residing within the PAR on both the Z and W chromosomes. The concurrent discovery of *Amh* and *Amhr2* as duplicate copies of their autosomal orthologs (see Guo et al., 2025 GigaScience, this issue) on the Z chromosome and confirmation here that they do not reside on the W, hints at a dosage-based mechanism of sex determination involving one or both of these genes. *Amh* is a gene and its receptor *AmhR2* are central to male differentiation in vertebrates and so are predisposed to recruitment as master sex determining genes on the sex chromosomes. This has occurred multiple times in fish with the enlistment of *Amh* or *AmhR2* to the Y chromosome (Li et al., 2015; Song et al., 2021; Nakamoto et al., 2021; Jeffries et al., 2022) or the involvement of *Amh* in the establishment of a *de novo* sex chromosome (Kamiya et al., 2012). In the frog *Rana temporaria*, the Y chromosome underwent a reciprocal translocation with an autosome fusing them into a single inherited neo-Y chromosome that included key sex genes *Dmrt1*, *Amh*, and *AmhR2* (Rodrigues et al., 2016). *Amh* is also implicated as the master sex determining gene in monotremes (Zhou et al., 2021). Our results indicate that sex determination in the dragon is likely involve more complex gene interactions, involving expression of the Z and autosomal copies of *Amh* and *AmhR2* and involving also *Nr5a1* which encodes transcription factor SF1 and has a foundational involvement in sex determination in vertebrates. The gene *Nr5a1*, although on the PAR as confirmed here, and with virtually identical copies on the Z and W chromosomes, yields substantially different Z and W transcriptional isoform composition (Zhang et al., 2022). This suggests that complex interactions between these genes and their resultant transcription factors and intermediaries, determines sex in the bearded dragon. This will be a fruitful area for future investigation.

This annotated assembly for the central bearded dragon was generated as part of the AusARG initiative of Bioplatforms Australia, to contribute to the suite of high-quality genomes available for Australian reptiles and amphibians as a national resource. The central bearded dragon is already widely used in research requiring genomic foundations, in large part because of the earlier publication of an assembly based on short read technologies (Georges et al., 2015). The central bearded dragon is an emerging model species (Ollonen et al., 2018) because of its high fecundity and short incubation, ease with which it adapts to captivity and a published genome, all considered key advantages accelerating its use (Infante et al., 2018). We anticipate that this new and vastly improved reference genome will serve to accelerate comparative genomics, developmental studies and evolutionary research on this and other species. As an exemplar of a well-studied oviparous taxon with sex reversal by temperature, the central bearded dragon reference assembly will provide a solid basis for genomic studies of the evolution of the genetic basis for reprogramming of sexual development under the influence of environmental temperature (Quinn et al., 2007; Holleley, et al., 2015; Castelli et al., 2021).

## Availability of Supporting Data

The supplementary file contains a description of all supplemental materials, which include tables showing software used in the preparation of this paper, outcomes of the sequencing on the four sequencing platforms used, and figures in support of statements on the quality of data. The authors affirm that all other data necessary for confirming the conclusions of the article are present within the article, figures, and tables. The annotated assembly can be accessed from NCBI as PviZW2.1 (Accession No., to be provided on acceptance) and all reads used in support of the assembly are lodged with the Short Read Archive. Accession numbers are provided in the main text and the Supplementary Tables (Tables S2–S6). High resolution versions of figures and custom scripts used to conduct the analyses are at https://github.com/kango2/ausarg/.

## Abbreviations

BAC: Bacterial Artificial Chromosome
BUSCO: Benchmarking Universal Single-Copy Orthologs
EBP: Earth BioGenome Project
HiC: High-throughput Chromosome Conformation Capture
HiFi: High Fidelity
L50: min number of contigs (or scaffolds) to add in length to 50% of assembly length
L90: min number of contigs (or scaffolds) to add in length to 90% of assembly length
LINE: Long Interspersed Nuclear Element
LTR: Long Terminal Repeat
N50: median (50th percentile) contig or scaffold length
N90: 90th percentile of contig or scaffold length
NCBI: The National Center for Biotechnology information
ONT: Oxford Nanopore Technologies
PacBio: Pacific Biosciences
PAR: pseudoautosomal region
PCR: polymerase chain reaction
Q20: Phred score of 20 corresponding to a 1% error rate
Q30: Phred score of 30 corresponding to a 0.01% error rate
rDNA: ribosomal DNA
RNAseq: RNA-sequencing
rRNA: ribosomal RNA
SINE: Short Interspersed Nuclear Element
snRNA: small nuclear RNA
T2T: telomere-to-telomere
tRNA: transfer RNA
UTR: Untranslated Region

## Author Contributions

All authors contributed to the writing and editing of drafts of this manuscript. In addition, A.G. was the AusARG project lead and responsible for coordinating the initial proposal and securing the funding; A.L.M-R. contributed to the development of assembly pipelines; D.S.B.D – prepared samples, constructed figures and contributed to the initial conceptual work; H.R.P. led the assembly and development of related workflows and pipelines; I.W.D. and J.H. provided oversight of the data generation and supervision of subsequent analysis; J.K.C. developed the annotation workflow and pipelines; N.C.L performed phased HiC analyses. N.C.L. and H.J. examined variation between the haplotypes and the reference haplotype for analysis of trends in heterozygosity. Z.A.C. undertook the rDNA annotation. H.R.P oversaw the data generation, associated quality control and the submission to NCBI; K.A. was responsible under the supervision of H.R.P for data curation and management, constructing the automated assembly and annotation workflows, for the manual curation of the assembly & analysis and post-assembly analysis; P.D.W. with H.R.P. provided oversight of the assembly and annotation, interpretation of the Z and W scaffolds. L.X., C.E.H., S.W. and X.Z. contributed to interpretation of the sex chromosome genes and the implications for future work. A.R-H. and L.M-G. conducted chromosome contact analysis.

## Acknowledgements

We acknowledge the provision of computing and data resources provided by the Australian BioCommons Leadership Share (ABLeS) program. This program is co-funded by Bioplatforms Australia (enabled by the National Collaborative Research Infrastructure Strategy, NCRIS) and the National Computational Infrastructure (NCI).

## Supplementary Materials

### List of Tables

**Table S1.** A list of software used for the analyses reported in this paper including version numbers and where it can be accessed.

**Table S2.** Summary statistics for the raw Illumina RNA sequence data used to assemble the transcriptome and for annotation.

**Table S3.** Summary statistics for the raw PacBio HiFi sequence data used for the assembly.

**Table S4.** Summary statistics for the raw Oxford Nanopore sequence data used for the assembly.

**Table S5.** Summary statistics for the HiC sequence data used to scaffold the assembly.

**Table S6.** Summary statistics for the raw Illumina DNA sequence data.

**Table S7.** Bacterial Artificial Chromosome (BAC) sequences mapped to the assembly scaffolds for the bearded dragon *Pogona vitticeps*.

**Table S8.** Satellite repeat units of the genome assembly for the bearded dragon *Pogona vitticeps* collapsed into 45 distinct classes based on sequence similarity.

**Table S9.** Summary of the copy number and percentage of the bearded dragon (*Pogona vitticeps*) genome covered by repeat elements

### List of Figures

**Figure S1.** Comparison of average read quality values (QV) versus read length for the two sequencing technologies: Oxford Nanopore Technologies (ONT) and PacBio HiFi.

**Figure S2.** HiC contact maps for Haplotype 2 showing an assembly mis-join in the YAHS assembly. (a) The original contact map showing the mis-join; (b) the resolved assembly with the mis-join was resolved manually.

**Figure S3.** Figure showing identification of putative centromeres for the six macrochromosomes and 10 microchromosomes of the bearded dragon Pogona vitticeps.

**Figure S4.** Size distribution of the repetitive elements that could not be identified.

**Figure S5**. Annotation of the mitochondrial genome of *Pogona vitticeps* assembled using *flye* and annotated using *mitoHiFi*.

### Custom Scripts

Custom scripts used to conduct the analyses are at https://github.com/kango2/ausarg/.

**Script 1**: *pacbiobam2fastx.sh* A custom script to remove any reads containing PacBio adapter sequences and convert the .bam files to FASTQ.

**Script 2**: *calculateGC.py* A custom script to calculate GC content in non-overlapping sliding windows of 10 Kbp.

**Script 3.** *processtrftelo.py* A script to identify regions >600 bp that contained conserved vertebrate telomeric repeat motif (TTAGGG).

**Script 4.** *ribocop.py* A script to search for consecutive alignments of 18S, 5.8S, and 28S to determine rDNA sequences.

### High Resolution Figures

**Figures 4 (a-c)**. A plot of the 16 longest scaffolds (corresponding to the number of chromosomes of the bearded dragon *Pogona vitticeps*.

**Figures 10 (a-c).** Sex chromosome analysis. a) For each sequencing technology, boxplots of read depth in 10 Kbp windows of macrochromosomes, microchromosomes, the PAR, and Z and W specific scaffolds.

---

**Table S1.**
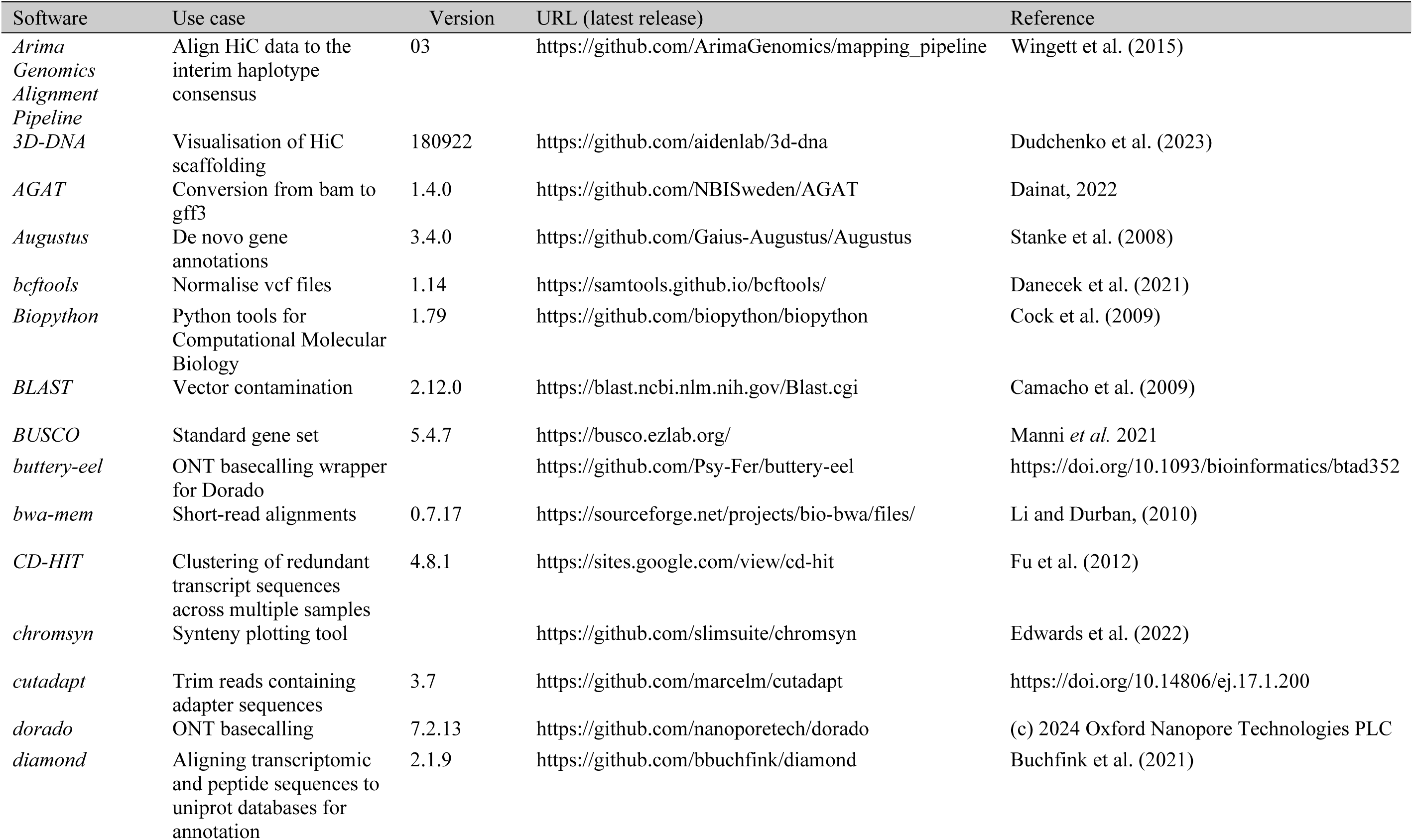
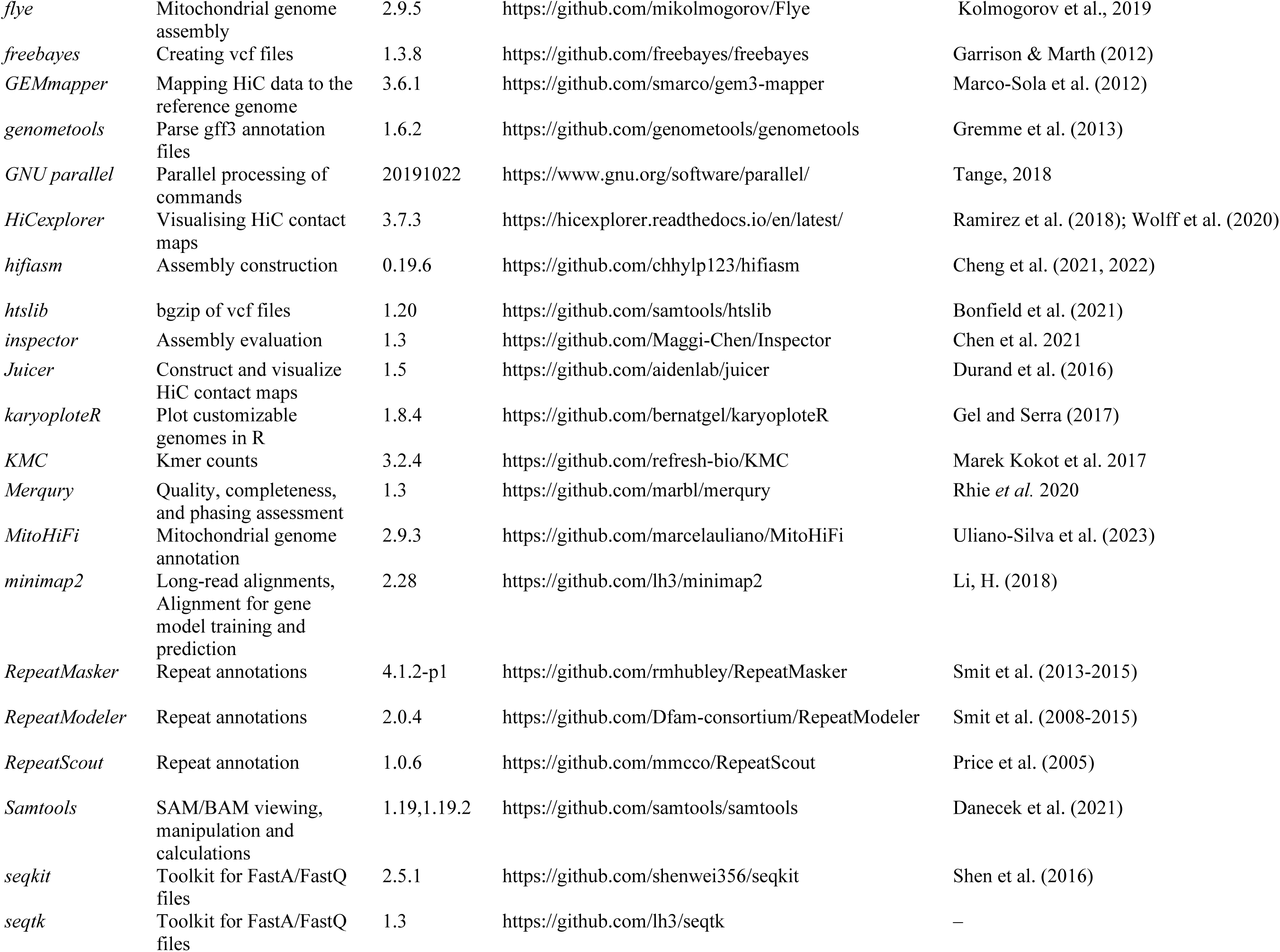
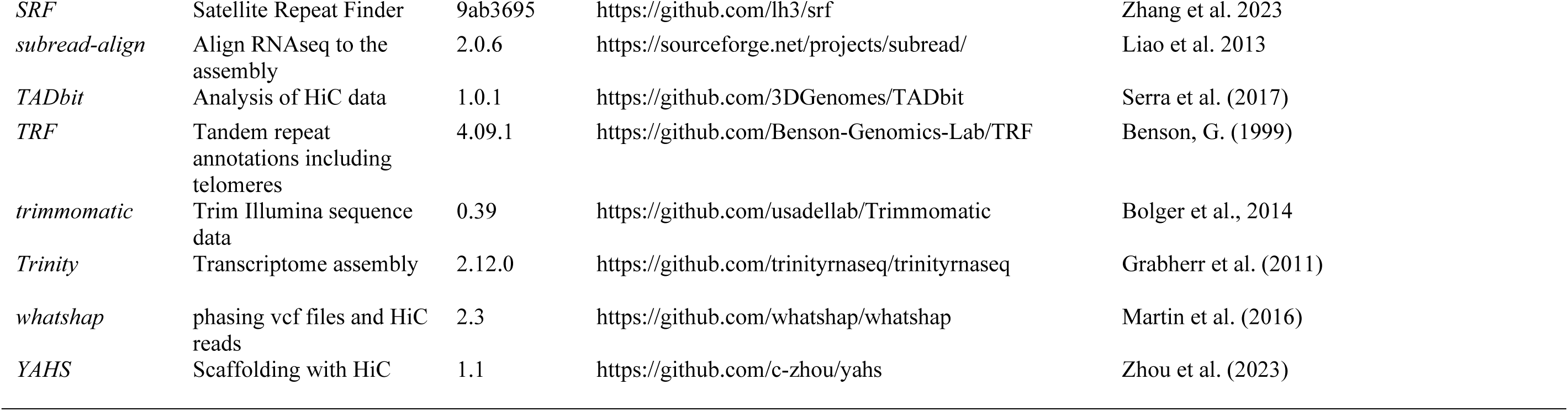
A list of software used for the analyses reported in this paper. Included are the use to which the software was put, the version number used, the source of the latest release of the software, and the associated published reference if available.

**Table S2.**
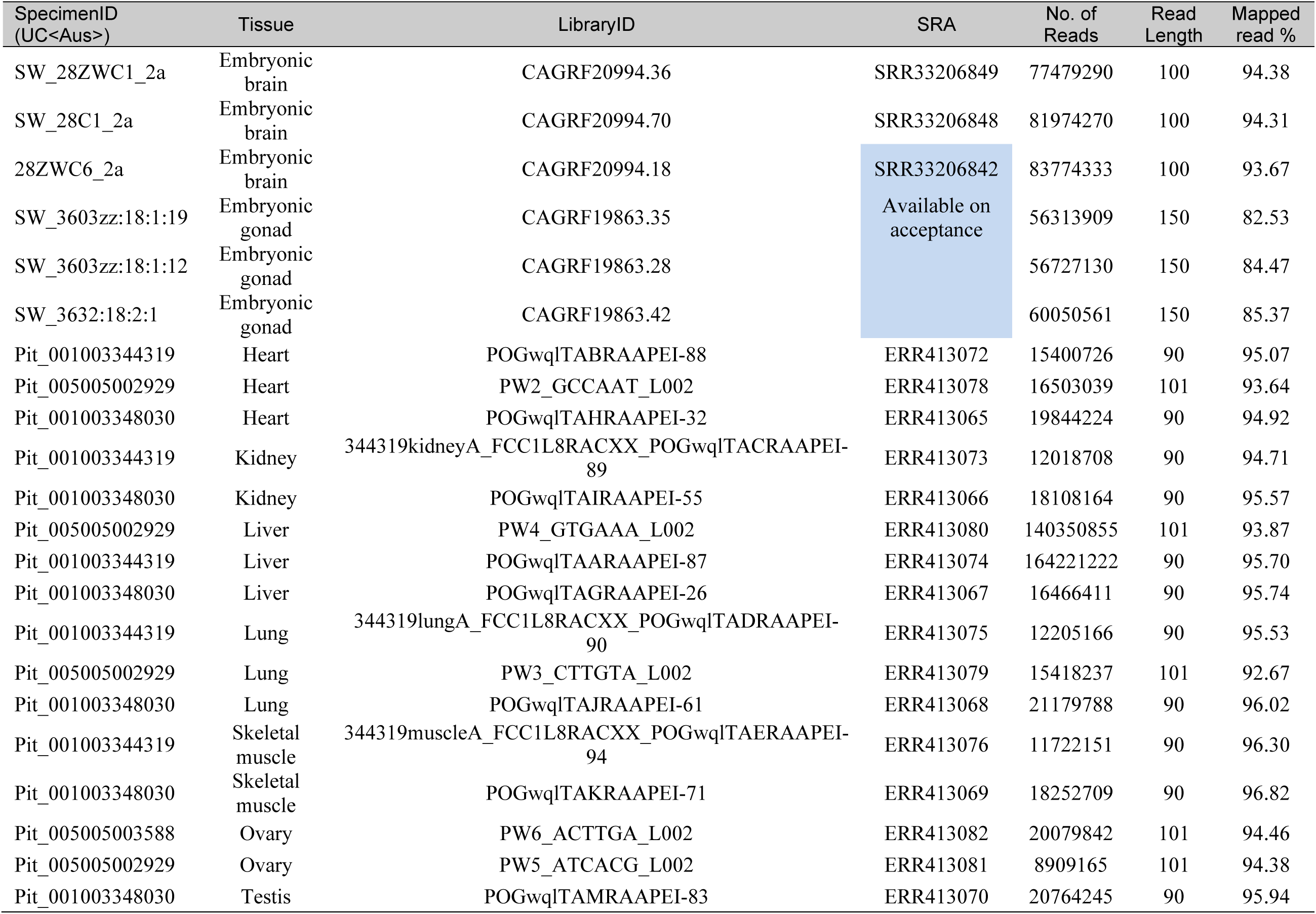
Summary statistics for the raw Illumina RNA sequence data used to assemble the transcriptome and for annotation.

**Table S3.** Summary statistics for the raw PacBio HiFi sequence data used for the assembly.

| SpecimenID<br>(UC<Aus>) | Tissue | Flow Cell | SRA | No. of Bases | No. of Reads | Read Length | N50 | N90 |
| --- | --- | --- | --- | --- | --- | --- | --- | --- |
| Pit_001003342236 | Blood | DAGPOG | SRR33206838 | 70,625,888,904 | 4,714,654 | 14,980 | 15,021 | 12,612 |

**Table S4.** Summary statistics for the raw Oxford Nanopore sequence data used for the assembly.

| SpecimenID<br>(UC<Aus>) | Tissue | Flow cell | SRA | No. of Bases | No. of Reads | Read Length<br>(bp) | N50 | N90 |
| --- | --- | --- | --- | --- | --- | --- | --- | --- |
| Pit_001003342236 | Blood | PAF09309 | SRR33206836 | 25,283,044,433 | 1,617,304 | 15,633 | 32,645 | 8,495 |
| Pit_001003342236 | Blood | PAF09661 | SRR33206837 | 39,491,153,126 | 3,394,876 | 11,633 | 32,142 | 5,155 |
| Pit_001003342236 | Blood | PAF10280 | SRR33206835 | 13,661,149,861 | 1,040,853 | 13,125 | 33,901 | 5,869 |
| Pit_001003342236 | Blood | PAF14969 | SRR33206845 | 3,934,999,270 | 332,739 | 11,826 | 34,339 | 4,907 |
| Pit_001003342236 | Blood | PAF21165 | SRR33206847 | 13,025,356,130 | 41,0291 | 31,747 | 59,981 | 16,099 |
| Pit_001003342236 | Blood | PAF32809 | SRR33206846 | 10,170,572,213 | 322,452 | 31,541 | 59,310 | 16,562 |

**Table S5.** Summary statistics for the HiC sequence data used to scaffold the assembly.

| SpecimenID<br>(UC<Aus>) | Tissue | Library | SRA | No. of Bases | No. of Reads | Read Length |
| --- | --- | --- | --- | --- | --- | --- |
| Pit_001003342236 | Blood | 350768_L001 | SRR33206844 | 22,349,129,631 | 148,007,481 | 151 |
| Pit_001003342236 | Blood | 350768_L002 | SRR33206843 | 22,248,518,784 | 14,7341,184 | 151 |

**Table S6.** Summary statistics for the Illumina sequence data.

| SpecimenID<br>(UC<Aus>) | Tissue | SRA | No. of Bases | No. of Reads | Mean Read Length |
| --- | --- | --- | --- | --- | --- |
| Pit_001003342236 | Blood | ERR409918, ERR409919, ERR409920 | 46,627,601,425 | 347,200,575 | 134 |

**Table S7.** Bacterial Artificial Chromosome (BAC) sequences mapped to the assembly scaffolds for the bearded dragon *Pogona vitticeps*. These BAC sequences were physically mapped to the chromosomes of the dragon by Deakin et al. (2016) and Young et al. (2013). They serve to confirm the association of the assembly scaffolds with the physical chromosomes. Scaffolds 1-6 correspond to Chromosomes 1-6, both assigned numbers by size. Scaffolds 7-15 correspond to microchromosomes numbered by Deakin et. al. (2016) as indicated. Scaffold 16, the pseudo-autosomal region of the Z and W sex chromosomes, is confirmed to correspond to the Z chromosome in the physical mapping of Clone 150H19. Scaffold 17 is confirmed to be associated with the W chromosome by the anchor Clone C1 of Quinn et al. (2010, Genbank EU938138).

| BAC ID | Chromosome | Scaffold | Start | End | Pairing | Length (bp) |
| --- | --- | --- | --- | --- | --- | --- |
| 16A1 | 1 | 1 | 123,407,376 | 123,573,280 | Double End | 165904 |
| 16A9 | 1 | 1 | 206,788,482 | 206,922,132 | Double End | 133650 |
| 57H2 | 1 | 1 | 23,855,452 | 23,978,560 | Double End | 123108 |
| 170F19 | 1 | 1 | 264,497,932 | 264,639,501 | Double End | 141569 |
| 184J20 | 1 | 1 | 273,284,146 | 273,381,615 | Double End | 97469 |
| 220D11 | 1 | 1 | 269,856,714 | 269,989,320 | Double End | 132606 |
| 220D7 | 1 | 1 | 198,454,349 | 198,556,275 | Double End | 101926 |
| 229E3 | 1 | 1 | 333,283,458 | 333,284,262 | Single End | 804 |
| 16A12 | 1 | 1 | 196,202,148 | 196,307,065 | Double End | 104917 |
| 11A17 | 1 | 1 | 157,690,828 | 157,788,565 | Double End | 97737 |
| 63H9 | 1 | 1 | 87,094,079 | 87,094,329 | Single End | 250 |
| 55I5 | 1 | 1 | 241,422,170 | 241,423,063 | Single End | 893 |
| 50I5 | 1 | 1 | 241,629,475 | 241,630,352 | Single End | 877 |
| 32C14 | 1 | 1 | 251,475,667 | 251,576,845 | Double End | 101178 |
| 31C8 | 1 | 1 | 159,024,102 | 159,024,242 | Single End | 140 |
| 240M14 | 1 | 1 | 219,470,320 | 219,470,616 | Single End | 296 |
| 191F7 | 1 | 1 | 302,271,442 | 302,349,405 | Double End | 77963 |
| 166M7 | 1 | 1 | 177,617,052 | 177,734,321 | Double End | 117269 |
| 161K22 | 1 | 1 | 312,800,053 | 312,800,696 | Single End | 643 |
| 12E8 | 1 | 1 | 275,882,655 | 275,883,474 | Single End | 819 |
| 57C5 | 1 | 1 | 113,655,489 | 113,656,282 | Single End | 793 |
| 28K11 | 1 | 1 | 233,289,187 | 233,289,570 | Single End | 383 |
| 231C17 | 1 | 1 | 139,946,455 | 139,946,593 | Single End | 138 |
| 167F2 | 1 | 1 | 189,870,499 | 189,871,006 | Single End | 507 |
| 16A23 | 2 | 2 | 66,988,935 | 67,151,660 | Double End | 162725 |
| 16A4 | 2 | 2 | 253,796,445 | 253,948,084 | Double End | 151639 |
| 176E5 | 2 | 2 | 211,772,680 | 211,923,226 | Double End | 150546 |
| 189J12 | 2 | 2 | 143,064,625 | 143,210,897 | Double End | 146272 |
| 195K1 | 2 | 2 | 166,137,002 | 166,292,352 | Double End | 155350 |
| 200H9 | 2 | 2 | 166,111,395 | 166,264,490 | Double End | 153095 |
| 219G15 | 2 | 2 | 39,640,903 | 39,810,574 | Double End | 169671 |
| 238E7 | 2 | 2 | 39,702,429 | 39,859,488 | Double End | 157059 |
| 203J2 | 2 | 2 | 146,656,877 | 146,757,171 | Double End | 100294 |
| 119M24 | 2 | 2 | 28,575,669 | 28,661,793 | Double End | 86124 |
| 76H7 | 2 | 2 | 301,486,841 | 301,597,372 | Double End | 110531 |
| 63C12 | 2 | 2 | 28,669,241 | 28,669,816 | Single End | 575 |
| 52J6 | 2 | 2 | 217,609,420 | 217,729,471 | Double End | 120051 |
| 21F11 | 2 | 2 | 93,857,250 | 93,857,797 | Single End | 547 |
| 213G6 | 2 | 2 | 139,763,286 | 139,763,386 | Single End | 100 |
| 160F4 | 2 | 2 | 29,206,132 | 29,283,406 | Double End | 77274 |
| 153E7 | 2 | 2 | 23,679,237 | 23,797,983 | Double End | 118746 |
| 74F13 | 2 | 2 | 77,615,637 | 77,615,778 | Single End | 141 |
| 57B1 | 2 | 2 | 123,317,551 | 123,318,392 | Single End | 841 |
| 42J16 | 2 | 2 | 203,443,365 | 203,443,969 | Single End | 604 |
| 225M3 | 2 | 2 | 4,920,160 | 4,920,229 | Single End | 69 |
| 104C4 | 2 | 2 | 31,514,680 | 31,515,011 | Single End | 331 |
| 185A1 | 3 | 3 | 228,674,547 | 228,798,733 | Double End | 124186 |
| 213B13 | 3 | 3 | 161,338,667 | 161,452,623 | Double End | 113956 |
| 214J17 | 3 | 3 | 154,656,138 | 154,797,939 | Double End | 141801 |
| 220D15 | 3 | 3 | 132,974,112 | 133,117,342 | Double End | 143230 |
| 221A23 | 3 | 3 | 72,300,526 | 72,406,473 | Double End | 105947 |
| 233A1 | 3 | 3 | 259,774,934 | 259,868,300 | Double End | 93366 |
| 71H17 | 3 | 3 | 212,191,524 | 212,191,947 | Single End | 423 |
| 60E8 | 3 | 3 | 101,809,810 | 101,809,872 | Single End | 62 |
| 41K6 | 3 | 3 | 263,235,206 | 263,235,636 | Single End | 430 |
| 224N13 | 3 | 3 | 54,477,440 | 54,572,033 | Double End | 94593 |
| 212O13 | 3 | 3 | 28,075,418 | 28,172,637 | Double End | 97219 |
| 98J10 | 3 | 3 | 48,936,383 | 49,023,996 | Double End | 87613 |
| 70H15 | 3 | 3 | 89,843,258 | 89,843,367 | Single End | 109 |
| 141N2 | 3 | 3 | 207,206,868 | 207,207,300 | Single End | 432 |
| 16A5 | 4 | 4 | 223,343,607 | 223,475,138 | Double End | 131531 |
| 219I19 | 4 | 4 | 15,956,020 | 16,099,868 | Double End | 143848 |
| 219N21 | 4 | 4 | 212,509,566 | 212,656,462 | Double End | 146896 |
| 230L10 | 4 | 4 | 30,902,447 | 31,039,140 | Double End | 136693 |
| 240P5 | 4 | 4 | 217,448,160 | 217,604,952 | Double End | 156792 |
| 94G20 | 4 | 4 | 131,828,670 | 131,828,800 | Single End | 130 |
| 55L2 | 4 | 4 | 35,486,665 | 35,486,806 | Single End | 141 |
| 39P7 | 4 | 4 | 22,140,696 | 22,140,795 | Single End | 99 |
| 240G13 | 4 | 4 | 70,024,998 | 70,025,094 | Single End | 96 |
| 6M2 | 4 | 4 | 46,157,770 | 46,158,228 | Single End | 458 |
| 47D22 | 4 | 4 | 50,742,605 | 50,742,749 | Single End | 144 |
| 43K4 | 4 | 4 | 197,338,787 | 197,338,848 | Single End | 61 |
| 152L3 | 4 | 4 | 44,449,933 | 44,450,480 | Single End | 547 |
| 16A22 | 5 | 5 | 21,793,409 | 21,907,879 | Double End | 114470 |
| 16A3 | 5 | 5 | 77,173,589 | 77,287,486 | Double End | 113897 |
| 210E16 | 5 | 5 | 191,209,567 | 191,327,937 | Double End | 118370 |
| 220D13 | 5 | 5 | 90,196,139 | 90,319,940 | Double End | 123801 |
| 233L23 | 5 | 5 | 191,071,018 | 191,295,551 | Double End | 224533 |
| 127M2 | 5 | 5 | 75,255,243 | 75,255,595 | Single End | 352 |
| 64B22 | 5 | 5 | 32,602,157 | 32,602,461 | Single End | 304 |
| 213P24 | 5 | 5 | 105,780,143 | 105,780,819 | Single End | 676 |
| 106M24 | 5 | 5 | 112,608,113 | 112,608,684 | Single End | 571 |
| 16A11 | 6 | 6 | 48,179,595 | 48,324,927 | Double End | 145332 |
| 132P11 | 6 | 6 | 98,533,797 | 98,652,352 | Double End | 118555 |
| 174P24 | 6 | 6 | 40,475,783 | 40,599,680 | Double End | 123897 |
| 200O10 | 6 | 6 | 115,445,513 | 115,564,715 | Double End | 119202 |
| 211I19 | 6 | 6 | 6,747,727 | 6,748,354 | Single End | 627 |
| 212P4 | 6 | 6 | 91,758,787 | 91,899,479 | Double End | 140692 |
| 225A2 | 6 | 6 | 113,902,256 | 114,060,808 | Double End | 158552 |
| 58O7 | 6 | 6 | 85,615,842 | 85,616,283 | Single End | 441 |
| 64E23 | 6 | 6 | 23,949,286 | 23,949,848 | Single End | 562 |
| 197P21 | 7 | 7 | 3,381,866 | 3,526,151 | Double End | 144285 |
| 75B13 | 7 | 7 | 4,323,835 | 4,324,297 | Single End | 462 |
| 163B7 | 7 | 7 | 7,515,270 | 7,515,386 | Single End | 116 |
| 55E1 | 7 | 7 | 27,269,115 | 27,269,479 | Single End | 364 |
| 220D8 | 11 | 8 | 19,977,289 | 20,074,547 | Double End | 97258 |
| 60I7 | Unplaced | 9 | 10,399,195 | 10,399,466 | Single End | 271 |
| 105P18 | 8 | 10 | 14,907,609 | 15,070,287 | Double End | 162678 |
| 232P19 | 8 | 10 | 3,325,360 | 3,419,255 | Double End | 93895 |
| 90K22 | 8 | 10 | 6,519,310 | 6,519,416 | Single End | 106 |
| 220D12 | 10 | 11 | 5,805,476 | 5,892,623 | Double End | 87147 |
| 161M1 | 10 | 11 | 25,752,319 | 25,752,921 | Single End | 602 |
| 188M22 | 11? | 12 | 3,335,567 | 3,499,827 | Double End | 164260 |
| 26E15 | 9 | 12 | 24,695,378 | 24,696,082 | Single End | 704 |
| 221B16 | 12 | 13 | 3,354,875 | 3,470,616 | Double End | 115741 |
| 39B11 | Unplaced | 13 | 9,469,199 | 9,469,292 | Single End | 93 |
| 214G3 | 14 | 14 | 13,829,067 | 13,993,455 | Double End | 164388 |
| 104H17 | Unknown<br>micro | 14 | 15,328,359 | 15,328,421 | Single End | 62 |
| 16A10 | 15 | 15 | 7,185,376 | 7,387,041 | Double End | 201665 |
| 185N3 | 13 | 15 | 2,286,971 | 2,409,964 | Double End | 122993 |
| 240C19 | 13 | 15 | 1,448,843 | 1,448,947 | Single End | 104 |
| 218G5 | Unknown<br>micro | 16 (PAR) | 3,169,270 | 3,169,542 | Single End | 272 |
| 150H19 | Z | 16 (PAR) | 3,169,270 | 3,169,542 | Single End | 272 |
| Contig C | Unplaced | 17 | 2,268,844 | 2,271,580 | -- | 2,736 |
| 57P12 | Unknown<br>micro | 18 | 2,381,218 | 2,381,349 | Single End | 131 |

**Table S8.** Satellite repeat units of the genome assembly for the bearded dragon *Pogona vitticeps* collapsed into 45 distinct classes based on sequence similarity.

| Satellite Repeat Name | Class | Repeat Unit Length (bp) |
| --- | --- | --- |
| POGVIT.v2.1#circ15-93 | srfclass-1 | 93 |
| POGVIT.v2.1#circ19-93 | srfclass-1 | 93 |
| POGVIT.v2.1#circ56-93 | srfclass-1 | 93 |
| POGVIT.v2.1#circ47-93 | srfclass-1 | 93 |
| POGVIT.v2.1#circ54-93 | srfclass-1 | 93 |
| POGVIT.v2.1#circ45-93 | srfclass-1 | 93 |
| POGVIT.v2.1#circ43-93 | srfclass-1 | 93 |
| POGVIT.v2.1#circ55-93 | srfclass-1 | 93 |
| POGVIT.v2.1#circ35-93 | srfclass-1 | 93 |
| POGVIT.v2.1#circ39-93 | srfclass-1 | 93 |
| POGVIT.v2.1#circ23-93 | srfclass-1 | 93 |
| POGVIT.v2.1#circ60-93 | srfclass-1 | 93 |
| POGVIT.v2.1#circ9-167 | srfclass-2 | 167 |
| POGVIT.v2.1#circ24-166 | srfclass-2 | 166 |
| POGVIT.v2.1#circ33-169 | srfclass-2 | 169 |
| POGVIT.v2.1#circ44-166 | srfclass-2 | 166 |
| POGVIT.v2.1#circ4-128 | srfclass-3 | 128 |
| POGVIT.v2.1#circ11-129 | srfclass-3 | 129 |
| POGVIT.v2.1#circ49-129 | srfclass-3 | 129 |
| POGVIT.v2.1#circ5-98 | srfclass-4 | 98 |
| POGVIT.v2.1#circ10-98 | srfclass-4 | 98 |
| POGVIT.v2.1#circ8-151 | srfclass-5 | 151 |
| POGVIT.v2.1#circ22-151 | srfclass-5 | 151 |
| POGVIT.v2.1#circ17-115 | srfclass-6 | 115 |
| POGVIT.v2.1#circ27-115 | srfclass-6 | 115 |
| POGVIT.v2.1#circ20-161 | srfclass-7 | 161 |
| POGVIT.v2.1#circ41-161 | srfclass-7 | 161 |
| POGVIT.v2.1#circ32-83 | srfclass-8 | 83 |
| POGVIT.v2.1#circ50-44 | srfclass-8 | 44 |
| POGVIT.v2.1#circ52-63 | srfclass-9 | 63 |
| POGVIT.v2.1#circ57-63 | srfclass-9 | 63 |
| POGVIT.v2.1#circ1-6 | srfclass-10 | 6 |
| POGVIT.v2.1#circ2-5695 | srfclass-11 | 5,695 |
| POGVIT.v2.1#circ3-151 | srfclass-12 | 151 |
| POGVIT.v2.1#circ6-150 | srfclass-13 | 150 |
| POGVIT.v2.1#circ7-151 | srfclass-14 | 151 |
| POGVIT.v2.1#circ12-398 | srfclass-15 | 398 |
| POGVIT.v2.1#circ13-157 | srfclass-16 | 157 |
| POGVIT.v2.1#circ14-230 | srfclass-17 | 230 |
| POGVIT.v2.1#circ16-9460 | srfclass-18 | 9,460 |
| POGVIT.v2.1#circ18-175 | srfclass-19 | 175 |
| POGVIT.v2.1#circ21-6 | srfclass-20 | 6 |
| POGVIT.v2.1#circ25-2190 | srfclass-21 | 2,190 |
| POGVIT.v2.1#circ26-115 | srfclass-22 | 115 |
| POGVIT.v2.1#circ28-249 | srfclass-23 | 249 |
| POGVIT.v2.1#circ29-179 | srfclass-24 | 179 |
| POGVIT.v2.1#circ30-236 | srfclass-25 | 236 |
| POGVIT.v2.1#circ31-5 | srfclass-26 | 5 |
| POGVIT.v2.1#circ34-144 | srfclass-27 | 144 |
| POGVIT.v2.1#circ36-150 | srfclass-28 | 150 |
| POGVIT.v2.1#circ37-102 | srfclass-29 | 102 |
| POGVIT.v2.1#circ38-3674 | srfclass-30 | 3,674 |
| POGVIT.v2.1#circ40-252 | srfclass-31 | 252 |
| POGVIT.v2.1#circ42-145 | srfclass-32 | 145 |
| POGVIT.v2.1#circ46-130 | srfclass-33 | 130 |
| POGVIT.v2.1#circ48-5 | srfclass-34 | 5 |
| POGVIT.v2.1#circ51-6 | srfclass-35 | 6 |
| POGVIT.v2.1#circ53-171 | srfclass-36 | 171 |
| POGVIT.v2.1#circ58-104 | srfclass-37 | 104 |
| POGVIT.v2.1#circ59-877 | srfclass-38 | 877 |
| POGVIT.v2.1#circ61-5 | srfclass-39 | 5 |
| POGVIT.v2.1#circ62-5 | srfclass-40 | 5 |
| POGVIT.v2.1#circ63-6 | srfclass-41 | 6 |
| POGVIT.v2.1#circ64-152 | srfclass-42 | 152 |
| POGVIT.v2.1#circ65-128 | srfclass-43 | 128 |
| POGVIT.v2.1#circ66-54 | srfclass-44 | 54 |
| POGVIT.v2.1#circ67-5 | srfclass-45 | 5 |

**Table S9.** Summary of the copy number and percentage of the bearded dragon (*Pogona vitticeps*) genome covered by repeat elements.

| Family | Numbers of elements | Length masked (bp) | % of sequence |
| --- | --- | --- | --- |
| **Retroelements** | **1,061,885** | **303,713,020** | **17.33** |
| SINEs | 114,787 | 15,769,671 | 0.90 |
| 5S | 6,625 | 553,278 | 0.03 |
| ID | 3,444 | 310,000 | 0.02 |
| MIR | 36,494 | 5,075,373 | 0.29 |
| U | 346 | 11,467 | 0.00 |
| U-L1 | 82 | 5,430 | 0.00 |
| tRNA | 15,532 | 1,687,441 | 0.10 |
| tRNA-Core-RTE | 8,936 | 319,829 | 0.02 |
| tRNA-Deu | 39,885 | 7,574,891 | 0.43 |
| tRNA-RTE | 3,443 | 231,962 | 0.01 |
| LINEs | 947,098 | 287,943,349 | 16.43 |
| CR1 | 178,263 | 39,822,603 | 2.27 |
| Dong-R4 | 21,177 | 6,300,007 | 0.36 |
| I | 649 | 44,872 | 0.00 |
| I-Jockey | 16,536 | 3,023,508 | 0.17 |
| L1 | 32,568 | 15,027,310 | 0.86 |
| L2 | 199,361 | 64,273,433 | 3.67 |
| Penelope | 66,521 | 13,708,273 | 0.78 |
| R2-NeSL | 116 | 22,059 | 0.00 |
| RTE-BovB | 372,010 | 129,982,167 | 7.42 |
| RTE-RTE | 269 | 64,349 | 0.00 |
| RTE-X | 50,133 | 11,827,799 | 0.67 |
| Rex-Babar | 9,495 | 3,846,969 | 0.22 |
| **LTR elements** | **80,454** | **72,706,738** | **4.15** |
| Copia | 3,713 | 3,224,584 | 0.18 |
| DIRS | 25,270 | 16,851,664 | 0.96 |
| ERV | 272 | 104,097 | 0.01 |
| ERV1 | 5,776 | 1,596,256 | 0.09 |
| ERVK | 1,795 | 996,283 | 0.06 |
| Gypsy | 27,479 | 42,285,692 | 2.41 |
| Ngaro | 16,149 | 7,648,162 | 0.44 |
| **DNA transposons** | **432,771** | **89,795,030** | **5.12** |
| CMC-Chapaev-3 | 385 | 154,910 | 0.01 |
| Maverick | 871 | 618,150 | 0.04 |
| PIF-Harbinger | 5,195 | 676,377 | 0.04 |
| TcMar | 58,667 | 18,056,702 | 1.03 |
| TcMar-Mariner | 2,350 | 1,083,990 | 0.06 |
| TcMar-Tc1 | 10,032 | 4,721,161 | 0.27 |
| TcMar-Tc2 | 120,600 | 15,583,345 | 0.89 |
| TcMar-Tigger | 120,825 | 27,876,875 | 1.59 |
| Zisupton | 1,384 | 316,647 | 0.02 |
| hAT-Ac | 24,713 | 4,868,609 | 0.28 |
| hAT-Blackjack | 6,949 | 1,247,319 | 0.07 |
| hAT-Charlie | 61,120 | 10,465,252 | 0.60 |
| hAT-Tag1 | 7,783 | 1,336,798 | 0.08 |
| hAT-Tip100 | 11,161 | 2,464,145 | 0.14 |
| hAT-hAT19 | 237 | 113,591 | 0.01 |
| hAT-hAT5 | 499 | 211,159 | 0.01 |
| **Penelope-like elements** | **1,797** | **150,905** | **0.01** |
| Chlamys | 1,797 | 150,905 | 0.01 |
| **Rolling-circles** | **447** | **95,044** | **0.01** |
| Helitron | 447 | 95,044 | 0.01 |
| **Unclassified** | **1,775,114** | **294,309,581** | **16.79** |
| **Total interspersed repeats** | **3,352,468** | **760,770,318** | **43.40** |
| Satellite | 4,246 | 1,284,141 | 0.07 |
| Simple Repeat | 576,326 | 34,276,116 | 1.96 |
| rRNA | 402 | 1,812,567 | 0.10 |
| snRNA | 2,397 | 502,447 | 0.03 |
| tRNA | 101 | 6,775 | 0.00 |
| **Total Masked** |  | **798,652,364** | **45.56** |

**Figure S1.**
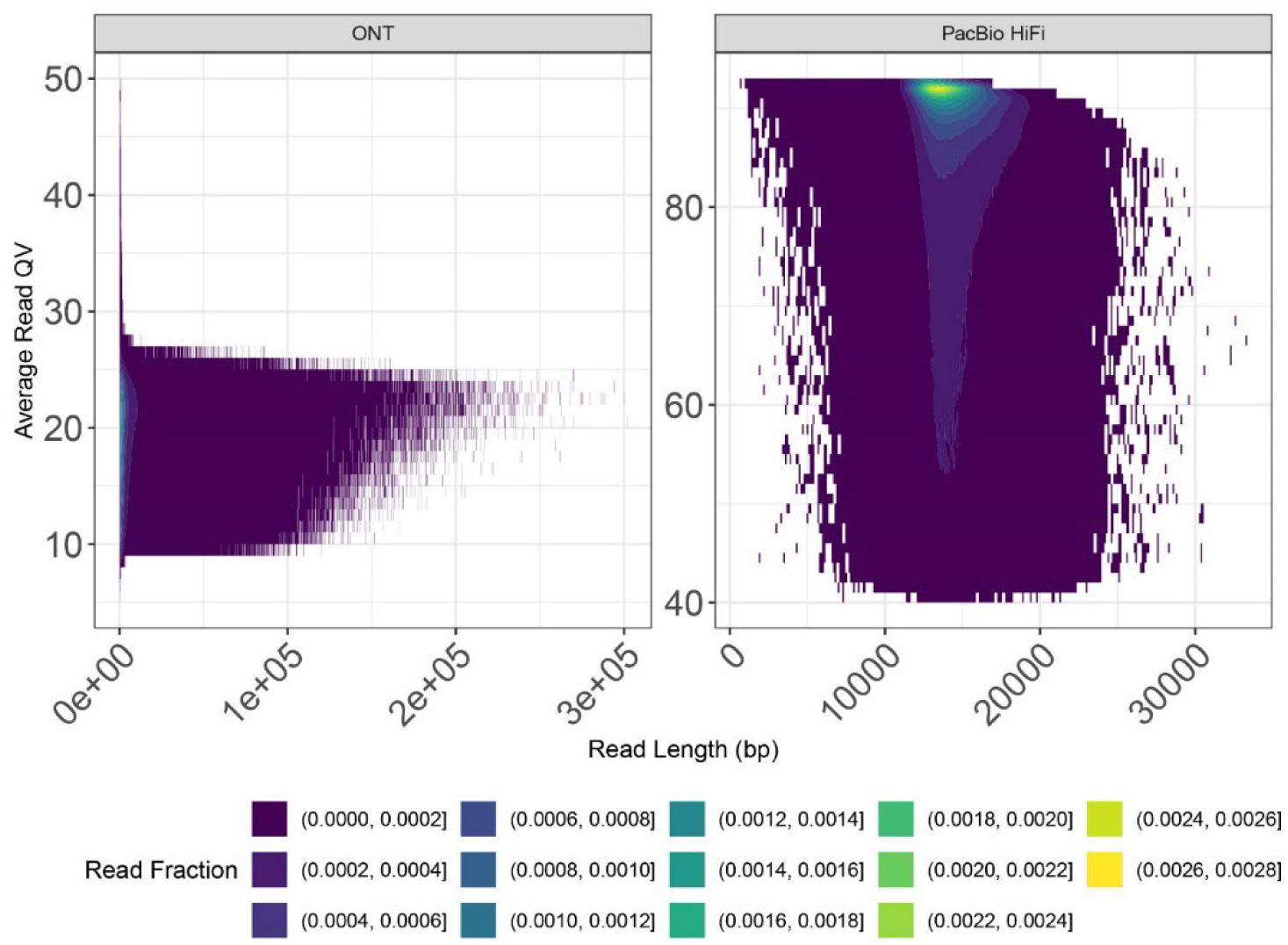
Comparison of average read quality values (QV) versus read length for the two sequencing technologies: Oxford Nanopore Technologies (ONT) and PacBio HiFi. Color intensity represents the base fraction in specified ranges, with darker colors indicating lower fractions and lighter colors indicating higher fractions. ONT reads show a broader distribution of read lengths with moderate quality values, whereas PacBio HiFi reads exhibit higher quality values with more concentrated reads at ~15 Kbp lengths.

**Figure S2.**
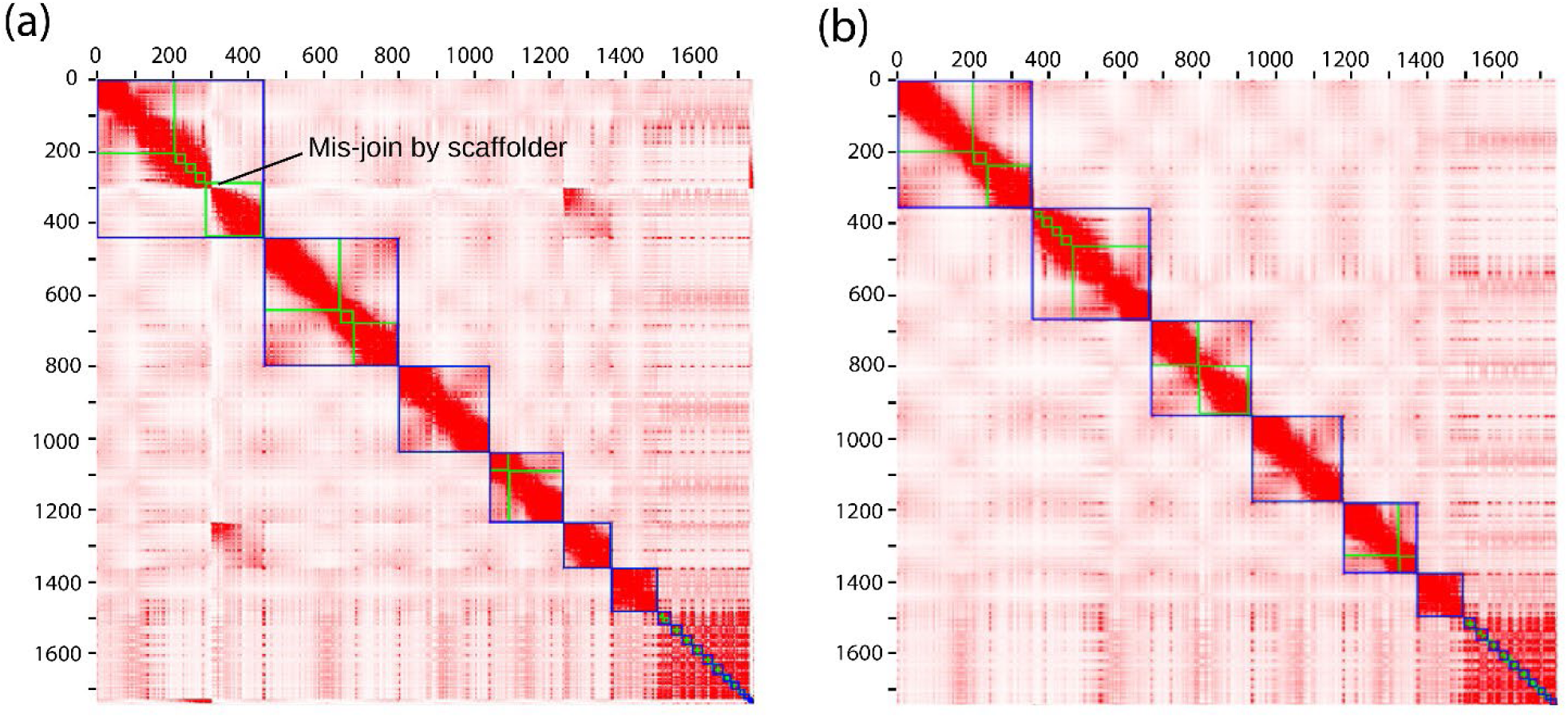
HiC contact maps for Haplotype 2 showing an assembly mis-join in the YAHS assembly. **(a)** The original contact map showing the mis-join; **(b)** the resolved assembly with the mis-join was resolved manually.

**Figure S3.**
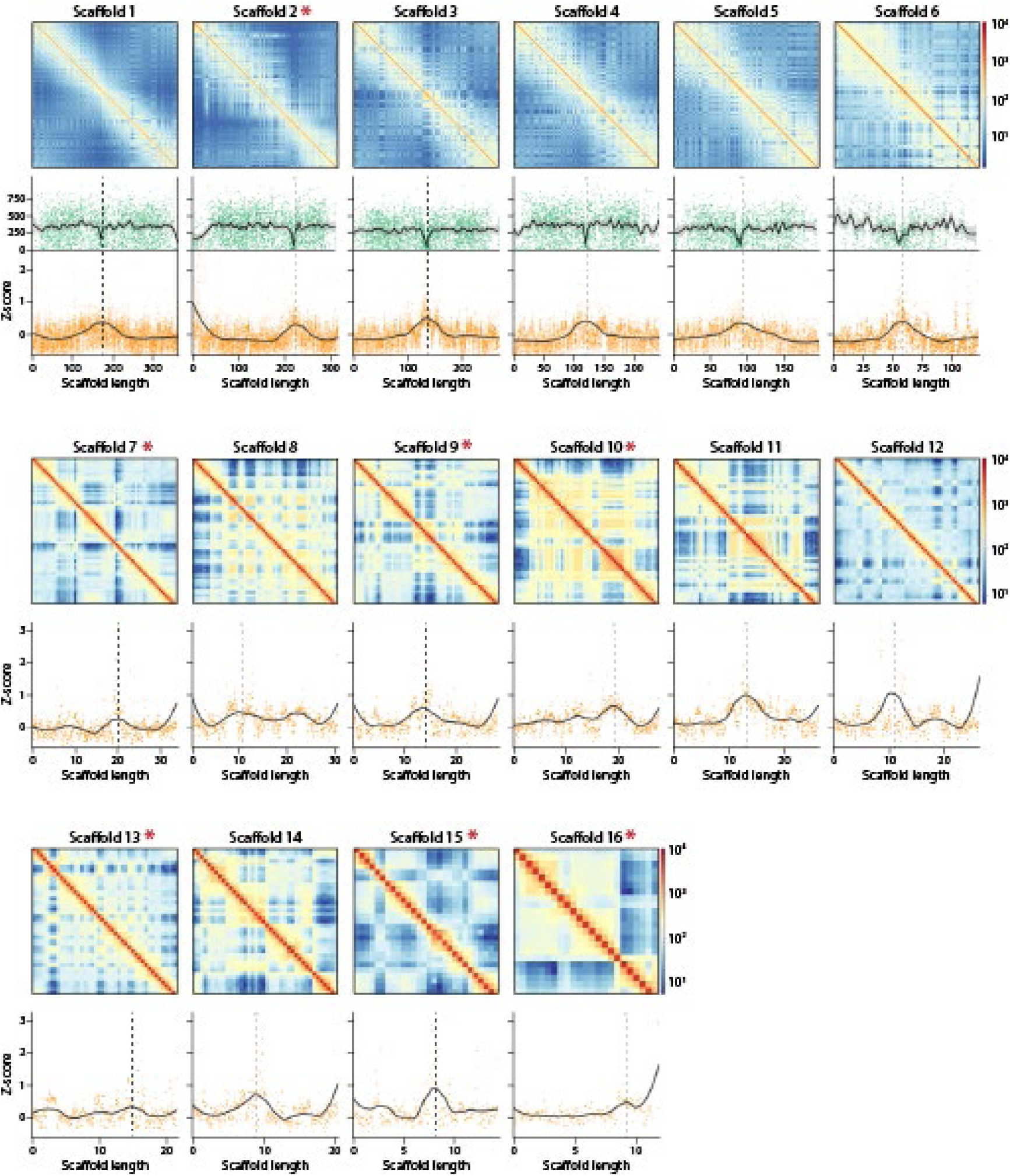
Identification of putative centromeres for the six macrochromosomes and 10 microchromosomes of the bearded dragon *Pogona vitticeps*. For both macrochromosomes and microchromosomes, the upper panels are chromosome-specific Hi-C heatmaps showing intra-chromosomal interactions. The lower panels are the Z-scores for the HiC inter-chromosomal interactions along chromosome length (Mbp) with smoothed lines of best fit. Each dot in the lower panels represents the Z-score interaction value of a different 50 Kbp bin. The middle panels for the macrochromosomes only, show the counts of heterozygous sites per 50 Kbp window (green dots) with lines of best fit and 95% confidence interval (grey shading). Dashed vertical lines correspond to putative centromere locations. Scaffold 16 is the pseudo-autosomal region (PAR) of the sex chromosomes. Scaffolds marked with an asterisk are inverted with respect to the published karyotype.

**Figure S4.**
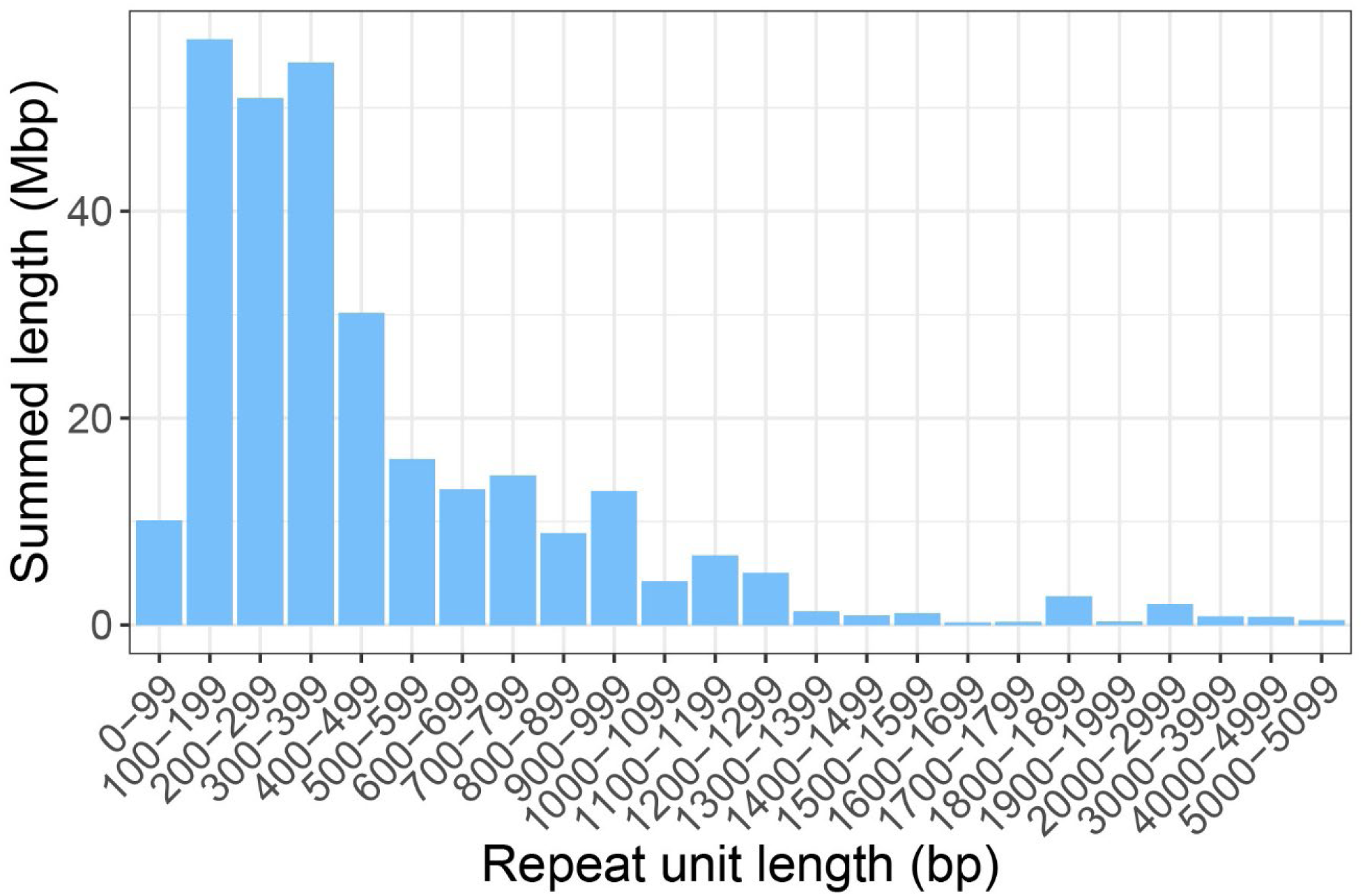
Size distribution of the repetitive elements that could not be identified (16.8%, Table S8).

**Figure S5.**
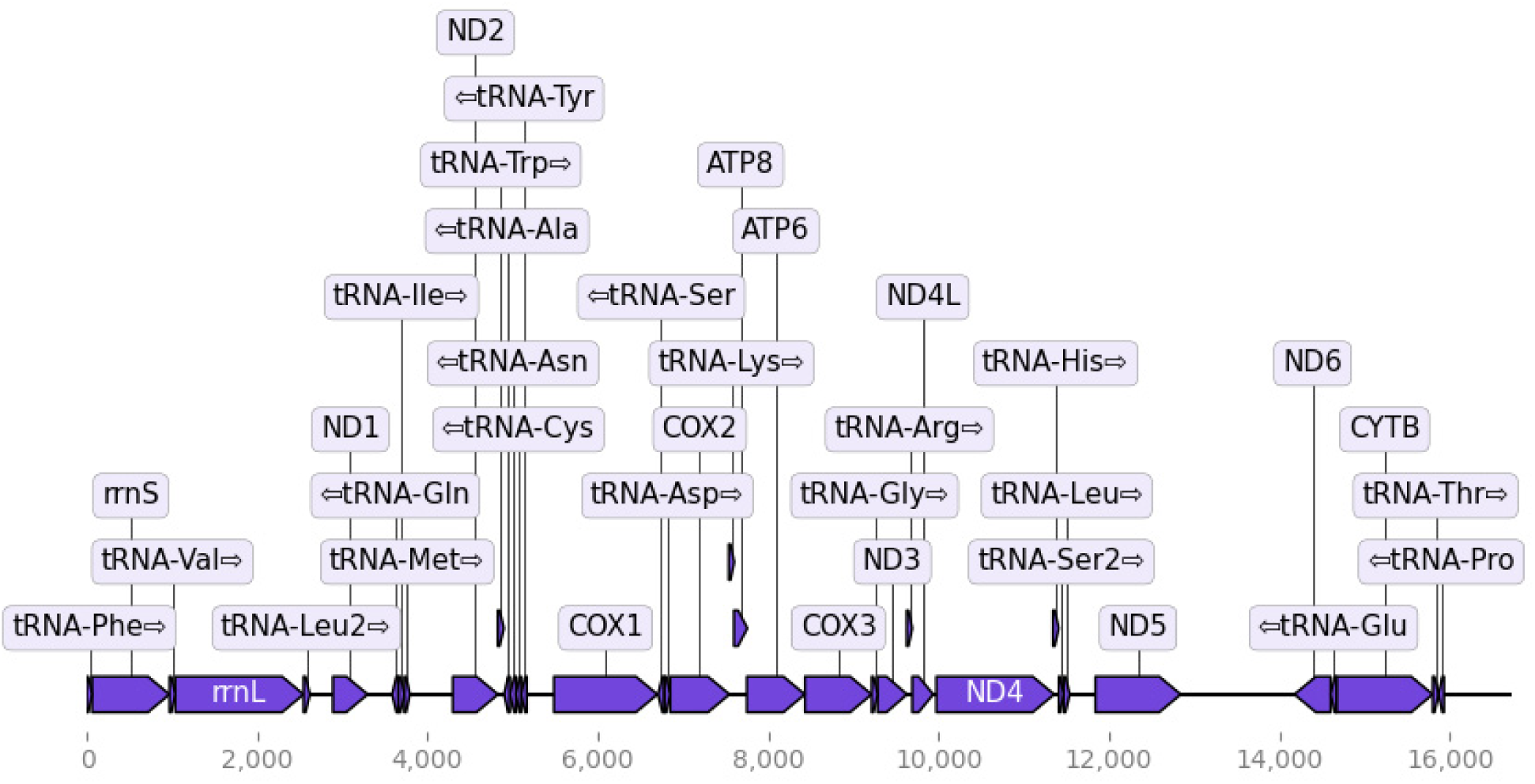
Annotation of the mitochondrial genome of the bearded dragon *Pogona vitticeps* assembled using *flye* and annotated using *mitoHiFi*. Control region not shown. Length 16,731 bp.

## References

1. Álvarez-González, L., Arias-Sardá, C., Montes-Espuña, L., Marín-Gual, L., Vara, C., Lister, N.C., Cuartero, Y., Garcia, F., Deakin, J., Renfree, M.B., Robinson, T.J., Martí-Renom, M.A., Waters, P.D., Farré, M., Ruiz-Herrera, A. 2022. Principles of 3D chromosome folding and evolutionary genome reshuffling in mammals. Cell Reports 41: 111839. doi: 10.1016/j.celrep.2022.111839.

2. Amer, S.A.M., Kumazawa, Y. 2005. Mitochondrial genome of *Pogona vitticepes* (Reptilia; Agamidae): control region duplication and the origin of Australasian agamids. Gene 346: 249–256. 10.1016/j.gene.2004.11.014.

3. Benson G. 1999. Tandem repeats finder: a program to analyze DNA sequences. Nucleic Acids Research 27:573–580. doi:10.1093/nar/27.2.573.

4. Bista, B., González-Rodelas, L., Álvarez-González, L., Wu, Z.Q, Montiel, E.E., Lee, L.S., Badenhorst, D.B., Radhakrishnan, S., Literman, R., Navarro-Dominguez, B., Iverson, J.B., Orozco-Arias, S., González, J., Ruiz-Herrera, A., Valenzuela, N. 2024. De novo genome assemblies of two cryptodiran turtles with ZZ/ZW and XX/XY sex chromosomes provide insights into patterns of genome reshuffling and uncover novel 3D genome folding in amniotes. Genome Research 34: 1553–1569. doi: 10.1101/gr.279443.124.

5. Bolger, A.M., Lohse, M., Usadel, B. 2014. Trimmomatic: a flexible trimmer for Illumina sequence data, Bioinformatics 30:2114–2120. 10.1093/bioinformatics/btu170.

6. Bonnan, M.F., Crisp, L.M., Barton, A., Dizinno, J., Muller, K., Smith, J., Walker, J. 2024. Exploring elbow kinematics in the central bearded dragon (*Pogona vitticeps*) using XROMM: Implications for the role of forearm long-axis rotation in non-avian reptile posture and mobility. The Anatomical Record. 10.1002/ar.25588.

7. Buchfink, B., Reuter, K., Drost, H.G. 2021. Sensitive protein alignments at tree-of-life scale using DIAMOND. Nature Methods 18:366–368. doi:10.1038/s41592-021-01101-x.

8. Chandrasekara, U., Mancuso, M., Sumner, J., Edwards, D., Zdenek, C.N., Fry, B.G. 2024. Sugar-coated survival: N-glycosylation as a unique bearded dragon venom resistance trait within Australian agamid lizards. Comparative Biochemistry and Physiology Part C: Toxicology & Pharmacology, 282,109929, 10.1016/j.cbpc.2024.109929.

9. Castelli, M., Georges, A., Cherryh, C., Rosauer, D., Sarre, S.D., Contador-Kelsall, I. and Holleley, C.E. 2021. Evolving thermal thresholds may explain the distribution of temperature sex reversal in an Australian dragon lizard (*Pogona vitticeps*). Diversity and Distributions 27:427–438.

10. Chen, Y., Zhang, Y., Wang, A.Y., Gao, M., Chong, Z. 2021. Accurate long-read de novo assembly evaluation with Inspector. Genome Biology 22:312. 10.1186/s13059-021-02527-4.

11. Cheng, H., Concepcion, G.T., Feng, X. et al. 2021. Haplotype-resolved de novo assembly using phased assembly graphs with hifiasm. Nature Methods 18, 170–175. 10.1038/s41592-020-01056-5.

12. Cheng, H., Jarvis, E.D., Fedrigo, O., Koepfli, K.P., Urban, L., Gemmell, N.J., Li, H. 2022. Haplotype-resolved assembly of diploid genomes without parental data. Nature Biotechnology 40:1332–1335. 10.1038/s41587-022-01261-x.

13. Cleveland, W.S. 1979. Robust Locally Weighted Regression and Smoothing Scatterplots. Journal of the American Statistical Association 74: 829–836.

14. Cogger, H.G. 2018. Reptiles and Amphibians of Australia (7^th^ updated ed.). Melbourne: CSIRO Publishing.

15. Camacho, C., Coulouris, G., Avagyan, V., Ma, N., Papadopoulos, J., Bealer, K., & Madden, T.L., 2009. BLAST+: architecture and applications. BMC Bioinformatics, 10, 421.

16. Danecek, P., Bonfield, J.K., Liddle, J., Marshall, J., Ohan, V., Pollard, M.O., Whitwham, A., Keane, T., McCarthy, S.A., Davies, R.M., Li, H. 2021. Twelve years of SAMtools and BCFtools. GigaScience 10, giab008. 10.1093/gigascience/giab008.

17. Deakin, J., Edwards, M.J., Patel, H., O’Meally, D., Lian, J., Stenhouse, R., Ryan, S., Livernois, A., Azad, B., Holleley, C., Li, Q. and Georges, A. 2016. Anchoring genome sequence to chromosomes of the central bearded dragon (*Pogona vitticeps*) enables reconstruction of ancestral squamate macrochromosomes and identifies sequence content of the Z chromosome. BMC Genomics 17:447.

18. Durand, N.C., Shamim, M.S., Machol, I., Rao, S.S.P., Huntley, M.H., Lander, E.S., Aiden, E.L. 2016. Juicer provides a one-click system for analyzing loop-resolution Hi-C experiments. Cell Systems 3: 95–98. 3(1):95-8. doi: 10.1016/j.cels.2016.07.002.

19. Edwards R.J., Dong C., Park R.F., Tobias P.A. 2022. A phased chromosome-level genome and full mitochondrial sequence for the dikaryotic myrtle rust pathogen, *Austropuccinia psidii”*. bioRxiv 2022.04.22.489119 doi: 10.1101/2022.04.22.489119.

20. Ehl, J., Altmanova, M., Kratochivil, L. 2021. With or without W? Molecular and cytogenetic markers are not sufficient for identification of environmentally-induced sex reversal in the bearded dragon. Sexual Development 15: 272–281. 10.1159/000514195.

21. Ezaz, T., Quinn, A.E., Miura, I., Sarre, S.D., Georges, A. and Graves, J.A.M. 2005. The dragon lizard *Pogona vitticeps* has ZZ/ZW micro-sex chromosomes. Chromosome Research 13:763–776.

22. Fenk, L.A., Riquelme, J.L., Laurent, G. 2024. Central pattern generator control of a vertebrate ultradian sleep rhythm. Nature 636:681–689. 10.1038/s41586-024-08162-w.

23. Fu, L., Niu, B., Zhu, Z., Wu, S., Li, W. 2013. CD-HIT: accelerated for clustering the next-generation sequencing data. Bioinformatics. 28:3150–152. doi: 10.1093/bioinformatics/bts565.

24. Georges, A., Li, Q., Lian, J., O’Meally, D., Deakin, J., Wang, Z., Zhang, P., Fujita, M., Patel, H.R., Holleley, C.E., Zhou, Y., Zhang, X., Matsurbara, K., Waters, P., Graves, J.A.M., Sarre, S.D. and Zhang, G. 2015. High-coverage sequencing and annotated assembly of the genome of the Australian dragon lizard *Pogona vitticeps*. GigaScience 4:45.

25. Grabherr, M.G., Haas, B.J., Yassour, M., Levin, J.Z., Thompson, D..A, Amit, I., Adiconis, X, Fan L., Raychowdhury, R., Zeng, Q., Chen, Z., Mauceli, E., Hacohen, N., Gnirke, A., Rhind, N., di Palma, F., Birren, B.W., Nusbaum, C., Lindblad-Toh, K., Friedman, N., Regev, A. 2011. Full-length transcriptome assembly from RNA-seq data without a reference genome. Nature Biotechnology 29:644–52. doi:10.1038/nbt.1883.

26. Gremme, G., Steinbiss, S., Kurtz, S. 2013. GenomeTools: a comprehensive software library for efficient processing of structured genome annotations. IEEE/ACM Trans Computational Biology and Bioinformatics10:645–656. doi: 10.1109/TCBB.2013.68.

27. Guo, Q., Pan, Y., Dai, W., Guo, F., Zeng, T., Chen, W., Mi, Y., Zhang, Y., Shi, S., Jiang, W., Cai, H., Wu, B., Zhou, Y., Wang, Y., Yang, C., Shi, X., Yan, X., Chen, J., Cai, C., Yang, J., Xu, X., Gu, Y., Dong, Li, Q. 2025. A near-complete genome assembly of the bearded dragon *Pogona vitticeps* provides insights into the origin of *Pogona* sex chromosomes. GigaScience, in press.

28. Holleley, C.E., O’Meally, D., Sarre, S.D., Graves, J.A.M., Ezaz, T., Matsubara, K., Azad, B., Zhang, X. and Georges, A. 2015. Sex reversal triggers the rapid transition from genetic to temperature-dependent sex. Nature 523:79–82.

29. Infante, C.R., Rasys, A.M., Menke, D.B. (2018). Appendages and gene regulatory networks: Lessons from the limbless. Genesis 56: e23078.

30. Jeffries D.L., Mee, J.A., Peichel, C.L. 2022. Identification of a candidate sex determination gene in *Culaea inconstans* suggests convergent recruitment of an *Amh* duplicate in two lineages of stickleback. Journal of Evolutionary Biology 35: 1683–1694.

31. Kamiya, T., Kai, W., Tasumi, S., Oka, A., Matsunaga, T., Mizuno, N., Fujita, M., Suetake, H., Suzuki, S., Hosoya, S., Tohari, S., Brenner, S., Miyadai, T., Venkatesh, B., Suzuki, Y., Kikuchi, K. 2012. A trans-species missense SNP in Amhr2 is associated with sex determination in the tiger pufferfish, *Takifugu rubripes* (fugu). PLoS Genetics 8:e1002798. doi: 10.1371/journal.pgen.1002798.

32. Kolmogorov, M., Yuan, J., Lin, Y., Pevzner, P. 2019. Assembly of long error-prone reads using repeat graphs. Nature Biotechnology 37:540–546. doi:10.1038/s41587-019-0072-8.

33. Lawniczak, M.K.N., Durbin, R., Flicek, P., +41, Richards, S. 2022. Standards recommendations for the Earth BioGenome Project. PNAS 119:e2115639118.

34. Li, H. 2018. Minimap2: pairwise alignment for nucleotide sequences, Bioinformatics 34:3094– 3100. 10.1093/bioinformatics/bty191.

35. Li, M., Sun, Y., Zhao, J., Shi, H., Zeng, S., Ye, K., Jiang, D., Zhou, L., Sun, L., Tao, W., Nagahama, Y., Kocher, T.D., Wang, D. 2015. A tandem duplicate of anti-Müllerian hormone with a missense SNP on the Y chromosome is essential for male sex determination in Nile Tilapia, *Oreochromis niloticus*. PLoS Genetics 11: e1005678.

36. Liao, Y., Smyth, G.K., Shi, W. 2013. The Subread aligner: fast, accurate and scalable read mapping by seed-and-vote. Nucleic Acids Research, 41:e108.

37. Macdonald, E., Whibley, A., Waters, P.D., Patel, H., Edwards, R.J., Ganley, A.R.D. 2024. Origin and maintenance of large ribosomal RNA gene repeat size in mammals. Genetics 228:iyae121, 10.1093/genetics/iyae121.

38. Manni, M., Berkeley, M.R., Seppey, M., Zdobnov, E.M. 2021. BUSCO: Assessing Genomic Data Quality and Beyond. Current Protocols 10.1002/cpz1.323.

39. Marco-Sola, S., Sammeth, M., Guigó, R., Ribeca, P. 2012. The GEM mapper: Fast, accurate and versatile alignment by filtration. Nature Methods 9:1185–1188. 10.1038/nmeth.2221

40. Miller S.A., Dykes DD, Polesky HF. A simple salting out procedure for extracting DNA from human nucleated cells. Nucleic Acids Res. 1988 Feb 11;16(3):1215. doi: 10.1093/nar/16.3.1215. PMID: 3344216; PMCID: PMC334765.

41. Mokhtaridoost, M., Chalmers, J.J., Soleimanpoor, M., McMurray, B.J., Lato, D.F., Nguyen, S.C., Musienko, V., Nash, J.O., Espeso-Gil, S., Ahmed, S., Delfosse, K., Browning, J.W.L., Barutcu, A.R., Wilson, M.D., Liehr, T., Shlien, S., Aref, A., Joyce, E.F., Weise, A., Maass, P.G. 2024. Inter-chromosomal contacts demarcate genome topology along a spatial gradient. Nature Communications 15: 9813. 10.1038/s41467-024-53983-y.

42. Nagashima S., Yamaguchi S.T., Zhou Z., Norimoto H. 2024. Transient cooling resets circadian rhythms of locomotor activity in lizards. Journal of Biological Rhythms 39:607–613. doi:10.1177/07487304241273190.

43. Nakamoto, M., Uchino, T., Koshimizu, E., Kuchiishi, Y., Sekiguchi, R., Wang, L., Sudo, R., Endo, M., Guiguen, Y., Schartl, M., Postlethwait, J.H., Sakamoto, T. 2021. A Y-linked anti-Müllerian hormone type-II receptor is the sex-determining gene in ayu, *Plecoglossus altivelis*. PLoS Genetics 17: e1009705. 10.1371/journal.pgen.1009705

44. Ollonen, J., da Silva, F.O., Mahlow, K. and Di-Poil, N. 2018. Skull development, ossification pattern, and adult shape in the emerging lizard model organism *Pogona vitticeps*: A comparative analysis with other Squamates. Frontiers in Physiology 2018:00278.

45. Pasquesi, G.I.M., Adams, R.H., Card, D.C., Schield, D.R., Corbin, A.B., Perry, B.W., Reyes-Velasco, J., Ruggiero, R.P., Vandewege, M.W., Shortt, J.A., Castoe, T.A. 2018. Squamate reptiles challenge paradigms of genomic repeat element evolution set by birds and mammals. Nat Commun 9:2774. 10.1038/s41467-018-05279-1.

46. Quast C., Pruesse E., Yilmaz P., Gerken J., Schweer T., Yarza P., Peplies J., Glöckner F.O. 2013. The SILVA ribosomal RNA gene database project: improved data processing and web-based tools. Nucleic Acids Research 41(D1):D590–D596.

47. Quinn, A.E., Georges, A., Sarre, S.D., Guarino, F., Ezaz, T., and Graves, J.A.M. 2007. Temperature sex reversal implies sex gene dosage in a reptile. Science 316:411.

48. Quinn, A.E., Ezaz, T., Sarre, S.D., Graves, J.A.M. and Georges, A. 2010. Extension, single-locus conversion and physical mapping of sex chromosome sequences identify the Z microchromosome and pseudo-autosomal region in a dragon lizard, *Pogona vitticeps*. Heredity 104:410–417.

49. Ramírez, F., Bhardwaj, V., Arrigoni, L., Lam, K. C., Grüning, B.A., Villaveces, J., Habermann, B., Akhtar, A., Manke, T. 2018. High-resolution TADs reveal DNA sequences underlying genome organization in flies. Nature Communications 9: 189. 10.1038/s41467-017-02525-w

50. Razmadze, D., Salomies, L., Di-Poï, N. 2024. Squamates as a model to understand key dental features of vertebrates. Developmental Biology 516:1–19. 10.1016/j.ydbio.2024.07.011.

51. Rhie, A., Walenz, B.P., Koren, S., Phillippy, A.M. 2020. Merqury: reference-free quality, completeness, and phasing assessment for genome assemblies. Genome Biology 21:245. 10.1186/s13059-020-02134-9.

52. Rodrigues, N., Vuille, Y., Brelsford, A., Merilä, J., Perrin, N. 2016. The genetic contribution to sex determination and number of sex chromosomes vary among populations of common frogs (*Rana temporaria*). Heredity 117: 25–32. 10.1038/hdy.2016.22.

53. Serra, F., Baù, D., Goodstadt, M., Castillo, D., Filion, G.J., Marti-Renom, M.A. 2017. Automatic analysis and 3D-modelling of Hi-C data using TADbit reveals structural features of the fly chromatin colors. PLoS Computational Biology 13: e1005665. 10.1371/journal.pcbi.1005665.

54. Smit, A..FA., Hubley, R. 2008-2015. RepeatModeler Open-1.0. <http://www.repeatmasker.org>.

55. Smit, A.F.A., Hubley, R., Green, P. 2013-2015. RepeatMasker Open-4.0. http://www.repeatmasker.org.

56. Song, W., Xie, Y., Sun, M., Li, X., Fitzpatrick, C.K., Vaux, F., O’Malley, K.G., Zhang, Q., Qi, J., He, Y. 2021. A duplicated *amh* is the master sex-determining gene for *Sebastes* rockfish in the Northwest Pacific. Open Biolology 11(7):210063. doi: 10.1098/rsob.210063.

57. Stanke, M., Morgenstern, B. 2005. AUGUSTUS: a web server for gene prediction in eukaryotes that allows user-defined constraints. Nucleic Acids Research 33:W465–7. doi:10.1093/nar/gki458.

58. Vasimuddin Md, Misra, S., Li, H., Aluru, S. 2019. Efficient Architecture-Aware Acceleration of BWA-MEM for Multicore Systems. IEEE Parallel and Distributed Processing Symposium (IPDPS), 2019. 10.1109/IPDPS.2019.00041

59. Wagner, S., Whiteley, S.L., Castelli, M., Patel, H.R., Deveson, I.W., Blackburn, J., Holleley, C.E., Marshall Graves, J.A. and Georges, A. 2023. Gene expression of male pathway genes *sox9* and *amh* during early sex differentiation in a reptile departs from the classical amniote model. BMC Genomics 24:243, 10.1186/s12864-023-09334-0.

60. Waters, P.D., Patel, H.R., Ruiz-Herrera, A., Álvarez-González, L., Lister, N.C., Simakov, O., Ezaz, T., Kaur, P., Frere, C., Grützner, F., Georges, A. and Marshall Graves, J.A. 2021. Microchromosomes are building blocks of bird, reptile and mammal chromosomes. Proceedings of the National Academy of Sciences USA 118(45): e2112494118.

61. Whiteley, S.L., Holleley, C.E., Blackburn, J., Deveson, I.W., Wagner, S., Graves, J.A.M., Georges, A. 2021. Two transcriptionally distinct pathways drive female development in a reptile with genetic sex determination and temperature induced sex reversal. PLoS Genetics 17:e1009465.

62. Whiteley, S.L., Holleley, C.E. and Georges, A. 2022. Developmental dynamics of sex reprogramming by high incubation temperatures in a dragon lizard. BMC Genomics 23:322.

63. Witten J.G. 1983. Some karyotypes of Australian agamids (Reptilia: Lacertilia). Australian Journal of Zoology 31:533–540.

64. Young, M.J., O’Meally, D., Sarre, S.D., Georges, A. and Ezaz, T. 2013. Molecular cytogenetic map of the central bearded dragon *Pogona vitticeps* (Squamata: Agamidae). Chromosome Research 21:361–374.

65. Zhang, X., Wagner, S., Deakin, J.E., Holleley, C.E., Matsubara, K., Deverson, I.W., Li, Z., Wang, C., O’Meally, D., Edwards, M., Patel, H.R., Ezaz, T., Marshall Graves, J.M. and Georges, A. 2022. Sex-specific splicing of Z- and W-borne nr5a1 alleles suggests sex determination is controlled by chromosome conformation PNAS (Proceedings of the National Academy of Sciences USA) 119(4):e2116475119.

66. Zhang Y, Chu J, Cheng H, Li H. 2023. De novo reconstruction of satellite repeat units from sequence data. Genome Research 33:1994–2001. doi: 10.1101/gr.278005.123. PMID: 37918962; PMCID: PMC10760446.

67. Zhou, C., McCarthy, S.A., Durbin, R. 2023. YaHS: yet another Hi-C scaffolding tool. Bioinformatics, 39, btac808.

68. Zhou, Y., Shearwin-Whyatt, L., Li, J., Song, Z., Hayakawa, T., Stevens, D., Fenelon, J.C., Peel, E., Cheng, Y., Pajpach, F., Bradley, N., Suzuki, H., Nikaido, M., Damas, J., Daish, T., Perry, T., Zhu, Z., Geng, Y., Rhie, A., Sims, Y., Wood, J., Haase, B., Mountcastle, J., Fedrigo, O., Li, Q., Yang, H., Wang, J., Johnston, S.D., Phillippy, A.M., Howe, K., Jarvis, E.D., Ryder, O.A., Kaessmann, H., Donnelly, P., Korlach, J., Lewin, H.A., Graves, J., Belov, K., Renfree, M.B., Grutzner, F., Zhou, Q., Zhang, G. 2021. Platypus and echidna genomes reveal mammalian biology and evolution. Nature 592: 756–762 10.1038/s41586-020-03039-0.

## References

69. Price, A.L., Jones, N.C., Pevzner, P.A. 2005. *De novo* identification of repeat families in large genomes, Bioinformatics 21:i351–i358. 10.1093/bioinformatics/bti1018.

70. Benson G. 1999. Tandem repeats finder: a program to analyze DNA sequences. Nucleic Acids Research 27:573–580. doi:10.1093/nar/27.2.573.

71. Bolger, A.M., Lohse, M., Usadel, B. 2014. Trimmomatic: a flexible trimmer for Illumina sequence data. Bioinformatics 30:2114–2120. 10.1093/bioinformatics/btu170.

72. Buchfink, B., Reuter, K., Drost, H.G. 2021. Sensitive protein alignments at tree-of-life scale using DIAMOND. Nature Methods 18:366–368. doi:10.1038/s41592-021-01101-x.

73. Camacho, C., Coulouris, G., Avagyan, V., Ma, N., Papadopoulos, J., Bealer, K., & Madden, T.L., 2009. BLAST+: architecture and applications. BMC Bioinformatics 10: 421.

74. Chen, Y., Zhang, Y., Wang, A.Y., Gao, M., Chong, Z. 2021. Accurate long-read de novo assembly evaluation with Inspector. Genome Biology 22:312. 10.1186/s13059-021-02527-4.

75. Cheng, H., Concepcion, G.T., Feng, X. et al. 2021. Haplotype-resolved de novo assembly using phased assembly graphs with hifiasm. Nature Methods 18, 170–175. 10.1038/s41592-020-01056-5.

76. Cheng, H., Jarvis, E.D., Fedrigo, O., Koepfli, K.P., Urban, L., Gemmell, N.J., Li, H. 2022. Haplotype-resolved assembly of diploid genomes without parental data. Nature Biotechnology 40:1332–1335. 10.1038/s41587-022-01261-x.

77. Cock, P.J.A., Antao, T., Chang, J.T., Chapman, B.A., Cox, C.J., Dalke, A., Friedberg, I., Hamelryck, T., Kauff, F., Wilczynski, B., de Hoon, M.J.L. 2009. Biopython: freely available Python tools for computational molecular biology and bioinformatics, Bioinformatics 25:1422–1423. 10.1093/bioinformatics/btp163

78. Danecek, P., Bonfield, J.K., Liddle, J., Marshall, J., Ohan, V., Pollard, M.O., Whitwham, A., Keane, T., McCarthy, S.A., Davies, R.M., Li, H. 2021. Twelve years of SAMtools and BCFtools. GigaScience 10, giab008. 10.1093/gigascience/giab008.

79. Dainat, J. 2022. Another Gtf/Gff Analysis Toolkit (AGAT): Resolve interoperability issues and accomplish more with your annotations. Plant and Animal Genome XXIX Conference. https://github.com/NBISweden/AGAT.

80. Dudchenko, O., Batra, S.S., Omer, A.D., Nyquist, S.K., Hoeger, M., Durand, N.C., Shamim, M.S., Machol, I., Lander, E.S., Aiden, A.P., Aiden, E.L. (2017). De novo assembly of the *Aedes aegypti* genome using Hi-C yields chromosome-length scaffolds. Science 356:92–95. doi: 10.1126/science.aal3327.

81. Durand, N.C., Shamim, M.S., Machol, I., Rao, S.S.P., Huntley, M.H., Lander, E.S., Aiden, E.L. 2016. Juicer provides a one-click system for analyzing loop-resolution Hi-C experiments. Cell Systems 3: 95–98. 3(1):95-8. doi: 10.1016/j.cels.2016.07.002.

82. Edwards R.J., Dong C., Park R.F., Tobias P.A. 2022. A phased chromosome-level genome and full mitochondrial sequence for the dikaryotic myrtle rust pathogen, Austropuccinia psidii”. bioRxiv 2022.04.22.489119 doi: 10.1101/2022.04.22.489119

83. Fu, L., Niu, B., Zhu, Z., Wu, S., Li, W. 2013. CD-HIT: accelerated for clustering the next-generation sequencing data. Bioinformatics. 28:3150–152. doi: 10.1093/bioinformatics/bts565.

84. Garrison E, Marth G. 2012. Haplotype-based variant detection from short-read sequencing. *arXiv preprint arXiv:1207.3907 [q-bio.GN]*.

85. Gel, B., Serra, E. 2017. karyoploteR: an R/Bioconductor package to plot customizable genomes displaying arbitrary data, Bioinformatics 33:3088–3090. 10.1093/bioinformatics/btx346.

86. Grabherr, M.G., Haas, B.J., Yassour, M., Levin, J.Z., Thompson, D..A, Amit, I., Adiconis, X, Fan L., Raychowdhury, R., Zeng, Q., Chen, Z., Mauceli, E., Hacohen, N., Gnirke, A., Rhind, N., di Palma, F., Birren, B.W., Nusbaum, C., Lindblad-Toh, K., Friedman, N., Regev, A. 2011. Full-length transcriptome assembly from RNA-seq data without a reference genome. Nature Biotechnology 29:644–52. doi:10.1038/nbt.1883.

87. Gremme, G., Steinbiss, S., Kurtz, S. 2013. GenomeTools: a comprehensive software library for efficient processing of structured genome annotations. IEEE/ACM Trans Computational Biology and Bioinformatics10:645–656. doi: 10.1109/TCBB.2013.68.

88. Kokot, M., Długosz, M., Deorowicz, S. 2017. KMC 3: counting and manipulating k-mer statistics, Bioinformatics 33:2759–2761. 10.1093/bioinformatics/btx304.

89. Olmogorov, M., Yuan, J., Lin, Y., Pevzner, P. 2019. Assembly of long error-prone reads using repeat graphs. Nature Biotechnology 37:540–546. doi:10.1038/s41587-019-0072-8.

90. Li, H. 2018. Minimap2: pairwise alignment for nucleotide sequences, Bioinformatics 34:3094–3100. 10.1093/bioinformatics/bty191

91. Liao, Y., Smyth, G.K., Shi, W. 2013. The Subread aligner: fast, accurate and scalable read mapping by seed-and-vote. Nucleic Acids Research, 41:e108.

92. Manni, M., Berkeley, M.R., Seppey, M., Zdobnov, E.M. 2021. BUSCO: Assessing Genomic Data Quality and Beyond. Current Protocols 10.1002/cpz1.323.

93. Marco-Sola, S., Sammeth, M., Guigó, R., Ribeca, P. 2012. The GEM mapper: Fast, accurate and versatile alignment by filtration. Nature Methods 9:1185–1188. 10.1038/nmeth.2221

94. Martin, M., Patterson, M., Garg, S., Fischer, S.O., Pisanti, N., Klau, G.W., Schoenhuth, A., Marschall, T. 2016. *WhatsHap:* fast and accurate read-based phasing. bioRxiv 085050. doi: 10.1101/085050

95. Price, A.L., Jones, N.C., Pevzner, P.A. 2005. *De novo* identification of repeat families in large genomes, Bioinformatics, Volume 21, Issue suppl_1, June 2005, Pages i351–i358, 10.1093/bioinformatics/bti1018.

96. Ramírez, F., Bhardwaj, V., Arrigoni, L., Lam, K. C., Grüning, B.A., Villaveces, J., Habermann, B., Akhtar, A., Manke, T. 2018. High-resolution TADs reveal DNA sequences underlying genome organization in flies. Nature Communications 9: 189. 10.1038/s41467-017-02525-w

97. Rhie, A., Walenz, B.P., Koren, S., Phillippy, A.M.. 2020. Merqury: reference-free quality, completeness, and phasing assessment for genome assemblies. Genome Biology 21:245. 10.1186/s13059-020-02134-9.

98. Serra, F., Baù, D., Goodstadt, M., Castillo, D., Filion, G.J., Marti-Renom, M.A. 2017. Automatic analysis and 3D-modelling of Hi-C data using TADbit reveals structural features of the fly chromatin colors. PLoS Computational Biology 13: e1005665. 10.1371/journal.pcbi.1005665.

99. Shen, W., Le, S., Li, Y., Hu, F. 2016. SeqKit: A cross-platform and ultrafast toolkit for FASTA/Q file manipulation. PLoS ONE 11(10): e0163962. 10.1371/journal.pone.0163962.

100. Smit, A.FA., Hubley, R. RepeatModeler Open-1.0. 2008-2015 <http://www.repeatmasker.org>.

101. Smit, A.F.A., Hubley, R., Green, P. RepeatMasker Open-4.0. 2013-2015 <http://www.repeatmasker.org>

102. Stanke, M., Morgenstern, B. 2005. AUGUSTUS: a web server for gene prediction in eukaryotes that allows user-defined constraints. Nucleic Acids Research 33:W465–7. doi:10.1093/nar/gki458.

103. Tange O. 2018. GNU Parallel 2018, March 2018, 10.5281/zenodo.1146014.

104. Uliano-Silva, M., Ferreira, J.G.R.N., Krasheninnikova, K., Darwin Tree of Life Consortium, Formenti, G., Abueg, L., Torrance, J., Myers, E.W., Durbin, R., Blaxter, M., McCarthy, S.A. 2023. MitoHiFi: a python pipeline for mitochondrial genome assembly from PacBio high fidelity reads. BMC Bioinformatics 24:288. 10.1186/s12859-023-05385-y.

105. Wingett, S., Ewels, P., Furlan-Magaril, M., Nagano, T., Schoenfelder, S., Fraser, P., Andrews, S. 2015. HiCUP: pipeline for mapping and processing Hi-C data. F1000Research 4:1310. doi: 10.12688/f1000research.7334.1

106. Wolff, J., Rabbani, L., Gilsbach, R., Richard, G., Manke, T., Backofen, R., Grüning, B.A. 2020. Galaxy HiCExplorer 3: a web server for reproducible Hi-C, capture Hi-C and single-cell Hi-C data analysis, quality control and visualization. Nucleic Acids Research 48:W177– W184. 10.1093/nar/gkaa220.

107. Zhang Y, Chu J, Cheng H, Li H. 2023. De novo reconstruction of satellite repeat units from sequence data. Genome Research 33:1994–2001. doi: 10.1101/gr.278005.123.

108. Zhou, C., McCarthy, S.A., Durbin, R. 2023. YaHS: yet another Hi-C scaffolding tool. Bioinformatics, 39, btac808

